# HPClas: A data-driven approach for identifying halophilic proteins based on catBoost

**DOI:** 10.1101/2023.11.30.569348

**Authors:** Shantong Hu, Xiaoyu Wang, Zhikang Wang, Yongfei Chang, Yun Tian, Zhuoqian Li, Menghan Jiang, Shihui Wang, Wenya Wang, Jiangning Song, Guimin Zhang

## Abstract

Halophilic proteins possess unique structural properties and exhibit high stability under extreme conditions. Such distinct characteristic makes them invaluable for applications in various aspects such as bioenergy, pharmaceuticals, environmental clean-up and energy production. Generally, halophilic proteins are discovered and characterized through labor-intensive and time-consuming wetlab experiments. Here, we introduced HPClas, a machine learning-based classifier developed using the catBoost ensemble learning technique to identify halophilic proteins. Extensive *in silico* calculations were conducted on a large public data set of 12574 samples and an independent test set of 200 sample pairs, on which HPClas achieved an AUROC of 0.877 and 0.845, respectively. The source code and curated data set of HPClas are publicly available at https://github.com/Showmake2/HPClas. In conclusion, HPClas can be explored as a promising tool to aid in the identification of halophilic proteins and accelerate their applications in different fields.

**Impact Statement:** In this study, we used a method based on prediction of proteins secreted by extreme halophilic bacteria to successfully extract a large number of halophilic proteins. Using this data, we have trained an accurate halophilic protein classifier that could determine whether an input protein is halophilic with a high accuracy of 84.5%. This research could not only promote the exploration and mining of halophilic proteins in nature, but also provide guidance for the generation of mutant halophilic enzymes.

## 1. Introduction

Since halophilic microorganism (halophiles) are adapted to extreme environments, they are conducive to pollution-free production that do not require sterilization(1). In addition, their proteins, which exhibit a strong dependency on high salt conditions(2), are referred to as halophilic proteins. Halophilic proteins commonly require high NaCl concentrations to be active, while halophilic enzymes depend on salt for their catalytic activity. For example, the α-amylase from *Halothermothnx orenii* exhibits high activity at 4.7 mol/L NaCl(3). Moreover, in-depth studies have found that some structural elements in halophilic enzymes, originally adapted for high salt conditions, can also help ensure their tolerance to organic solvents (4), which have wide applications in anhydrous environments. Moreover, halophilic enzymes also exhibit broad substrate specificity. These unique properties render them potentially valuable in various fields such as biofuel production, textile processing, waste treatment, and detergent additives (5). Halophilic proteins can also serve as crucial enzymes in building metabolic pathways throughout the production process(1).

Given the wide range of potential applications of halophilic proteins, their identification has become a major area of research interest. Many studies have collected samples of halophilic bacteria and archaea from salt lakes and oceans worldwide and employed whole-genome sequencing and wet laboratory validation methods to screen for halophilic proteins. Indeed, advances in DNA sequencing technologies, particularly the utilization of genomics and metagenomics tools, have facilitated the discovery of large numbers of protein sequences from a diverse spectrum of organisms. In 2014, Sharma et al. established a halophilic protein database HProtDB (6), and researchers noticed that most of the proteins contained in this database come from halophilic bacteria. However, proteins derived from halophilic bacteria may not be halophilic proteins. Some halophilic bacteria, such as *Halobacterium*, can effectively regulate their intracellular salt concentrations to be lower than the extracellular environment (7). They can also synthesize and accumulate compounds known as compatible solutes (e.g., proline, lysine, ectoine) to balance osmotic pressure inside and outside the cell. Therefore, these bacteria may contain some non-halophilic proteins. The current paradigm for halophilic protein identification relies on laboratory experiments to assess the halophilic characteristics of proteins. Nevertheless, this approach is quite inefficient and costly, and cannot meet the growing demand. Therefore, computational methods for halophilic protein screening are expected to address this issue and have attracted increasing research attention. However, previous methods suffered from severe data scarcity and performance generalization issues (8, 9). Furthermore, the lack of user-friendly software or websites hinders direct access by the public. Hence, there is a need for a powerful tool with greater accuracy that is readily available to the public to help unlock the potential of halophilic proteins.

In this study, we introduced a novel data collection method and created a large and comprehensive data set of halophilic proteins. On this basis, we reported a catBoost-based machine learning (ML) model termed HPClas (Halophilic Protein Classifier) for the identification of halophilic proteins (**Fig. 1**). HPClas was trained using high-quality data from UniProtKB and NCBI protein database, taking amino acid sequences as input and outputting probabilities of halophilic proteins. We compared the performance of HPClas with state-of-the-art (SOTA) models on the test set of a previous study. Experimental results demonstrated that HPClas outperformed existing halophilic protein prediction tools. Furthermore, we also conducted an interpretation analysis of our model, which further reinforced the credibility of our predictions.

**Figure 1.**
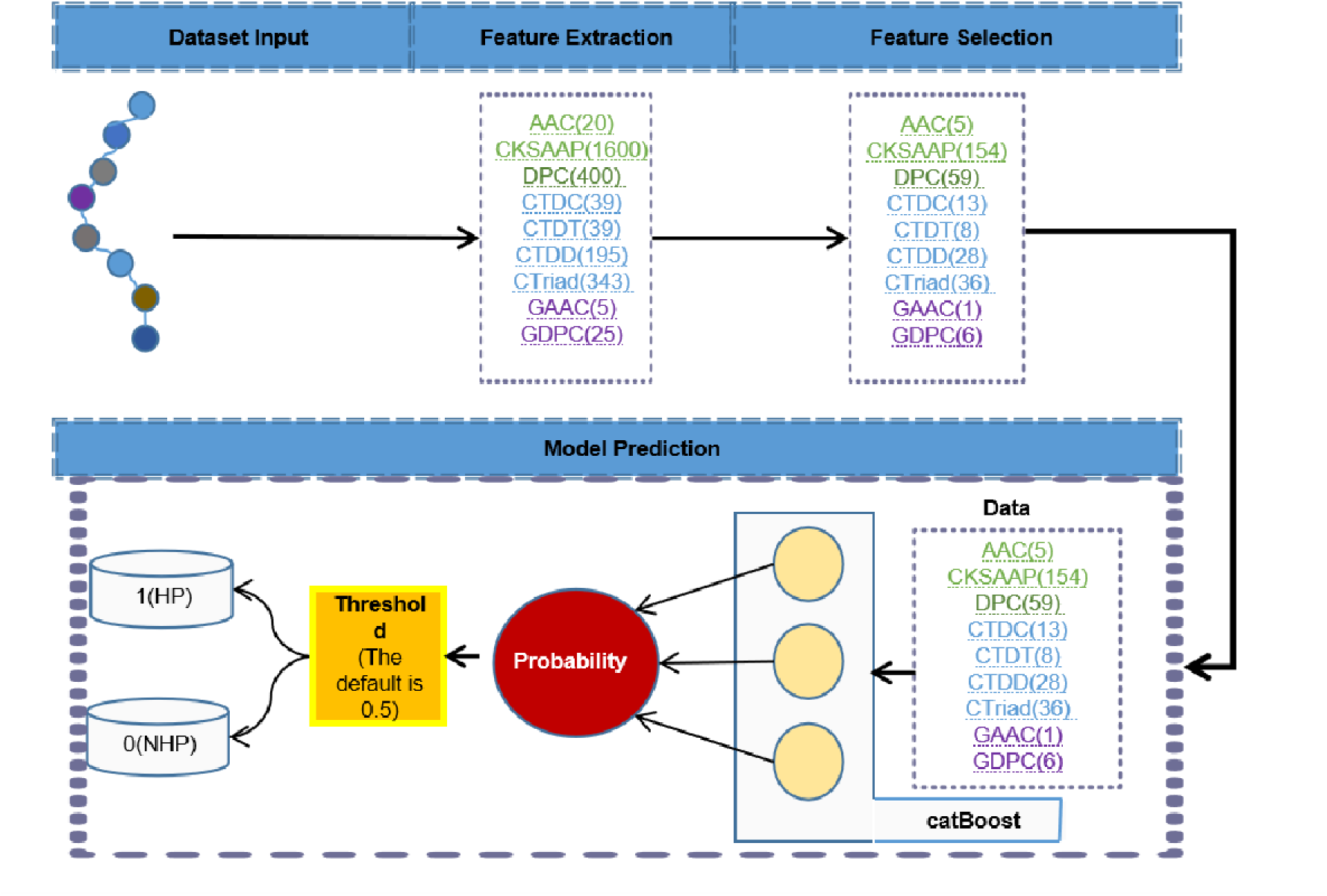
The overall architecture of HPClas. The input to HPClas is a protein sequence without a signal peptide. In the feature extraction process, features were extracted through feature descriptors, and the chi2 (10) was adopted for feature selection. 1 (HP) and 0 (NHP) represented halophilic protein and non-halophilic protein, respectively.

## 2. Results and discussion

### 2.1 Collection of a Large Training Dataset of Halophilic and Non-Halophilic proteins

In previous studies (8, 9), halophilic and non-haolphilic protein datasets were constructed but not made public, but all proteins in the genome of *Salinibacter ruber* DSM 13855 have been proved to be halophilic, whereas all proteins in the *Pelodictyo luteolum* DSM 2379 have been confirmed to be non-halophilic. We downloaded these proteins from UniProtKB (10) as part of the training dataset (Table 1).

**Table 1.**
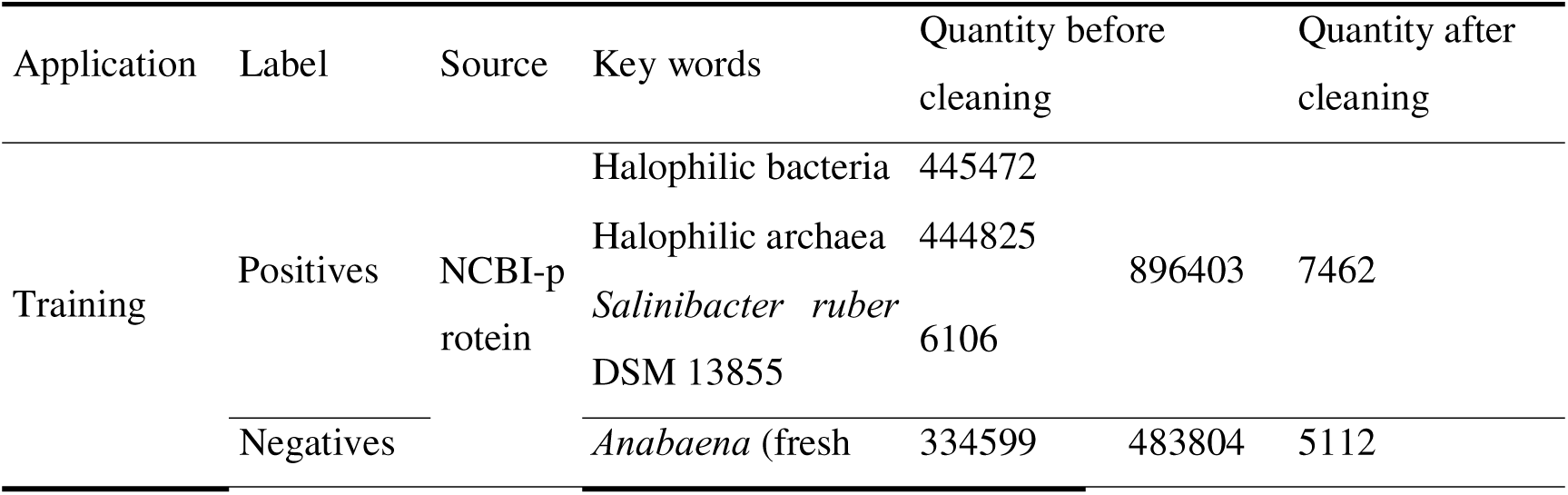
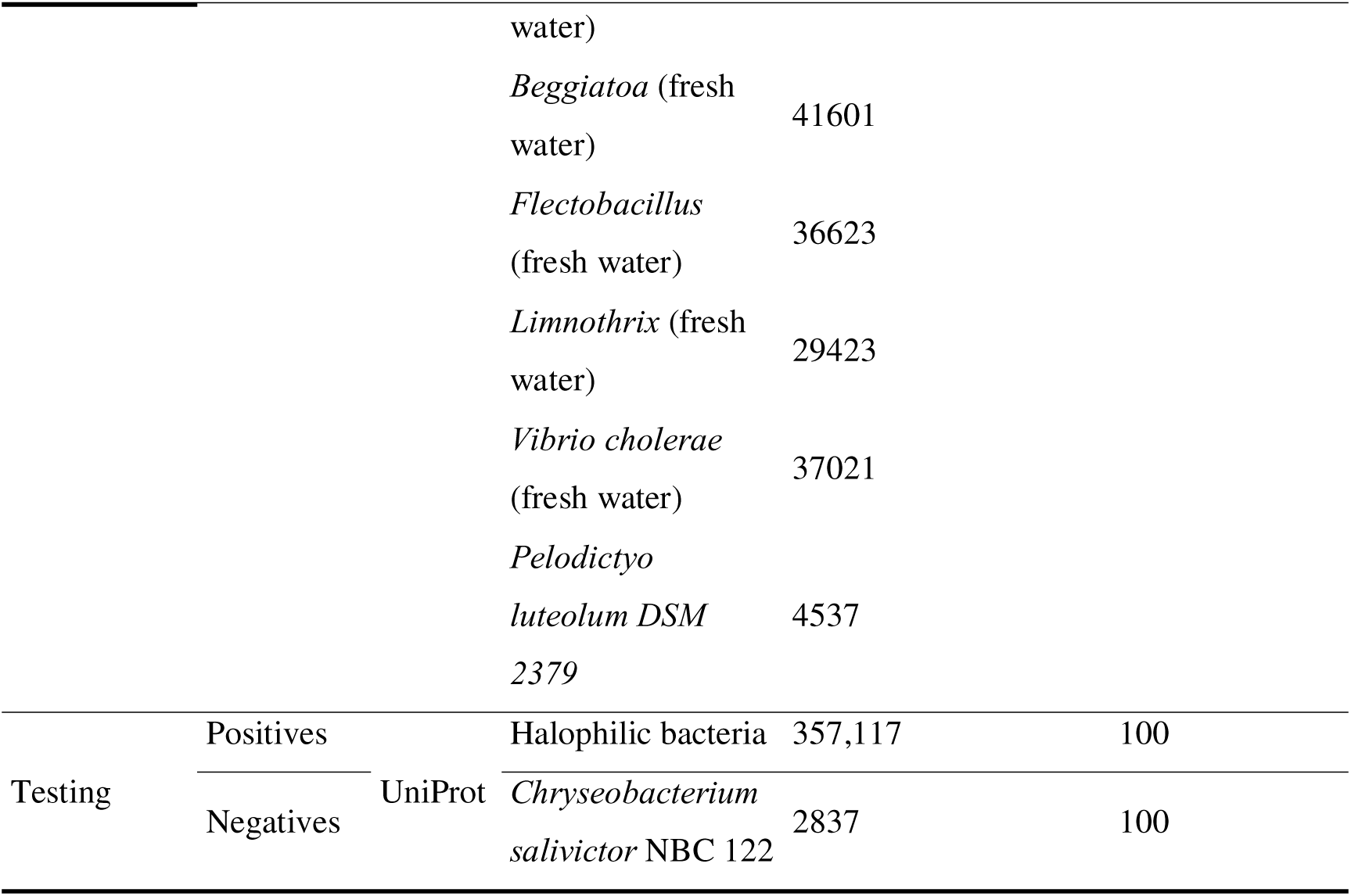
The overview of the training data set and independent test.

Additionally, a novel method here was proposed to generate a more comprehensive dataset of halophilic proteins. This method involves predicting signal peptides in proteins from halophilic microorganisms, enabling the identification of secreted proteins. These proteins are expected to adapt to high-salt conditions, thus categorizing them as halophilic proteins (11). The comprehensive data set contains training set and independent test set from the NCBI Protein Database and UniProtKB. Detailed information such as the application, labels, sources, key words, quantities before and after cleaning of the data set are summarized in **Table 1**. The training set contains 7462 halophilic proteins and 5112 non-halophilic proteins, while the test set contains 100 pairs of halophilic and non-halophilic proteins.

Wet experimentally validated halophilic proteins are extremely limited and challenging to retrieve, making the collection of a comprehensive data set quite difficult. However, Zhang et al (8, 9) have developed two classification models for halophilic proteins. For both models, the positive samples were from the halophilic bacterium, *S. ruber* DSM 13,855 (12), which did not rely on proton pumps to regulate salt concentration, ensuring that all proteins from this bacterium were halophilic proteins. Based on previous studies, we proposed a method to predict halophilic proteins by screening secreted proteins in halophilic bacteria. Secreted proteins that evolved in high-salt environments often function extracellularly, and tend to show a preference for salt tolerance. This strategy ensured the high quality and quantity of the current data set, while strict steps for removing redundancy including the removal of highly homologous sequences and the imposition of length criteria further reduce the risk of data leakage and ensured data quality. Moreover, proteins from multiple bacterial species increased the diversity of the data compared to proteins from a single strain source.

During the data mining process, not only did the internal homogeneity within the independent test set fall below 40%, but also the homogeneity between the entire independent test set and the training set was less than 25% to ensure a substantial distributional difference from the training set. Such high-quality data facilitates further assessing the model’s generalization, and thus, we chose it as a benchmark for optimizing the model.

Of course, our data have certain limitations. For example, the data sources are mainly secreted proteins, which may lead to potential misjudgments when predicting cytoplasmic proteins. To alleviate these limitations, we try to enrich the training dataset. For instance, the entire proteome of strain *S. ruber* DSM 13855 was used as a positive sample, and strain *P. luteolum* DSM 2379 as a negative sample. We hope to have more experimental data in the future that will allow us to obtain more intracellular protein data, so that we can further optimize our model.

### 2.2 Combination Screening of Models and Feature Encoding Methods

Eleven feature descriptors were employed to encode protein sequences into biologically meaningful vector representations, where each descriptor represents a statistical perspective on amino acid sequences. After feature extraction, we found that some feature values varied a lot, with some ranging from 0 to 0.01 and some from 1 to 1,000. These non-normalized features might considerably hinder model converge and affect network performance. Therefore, we employed the MinMaxScaler method to normalize the values of different features, ensuring that all values fell within the range of 0 to 1. The results are presented at https://github.com/Showmake2/HPClas/features.

**Figure 2.**
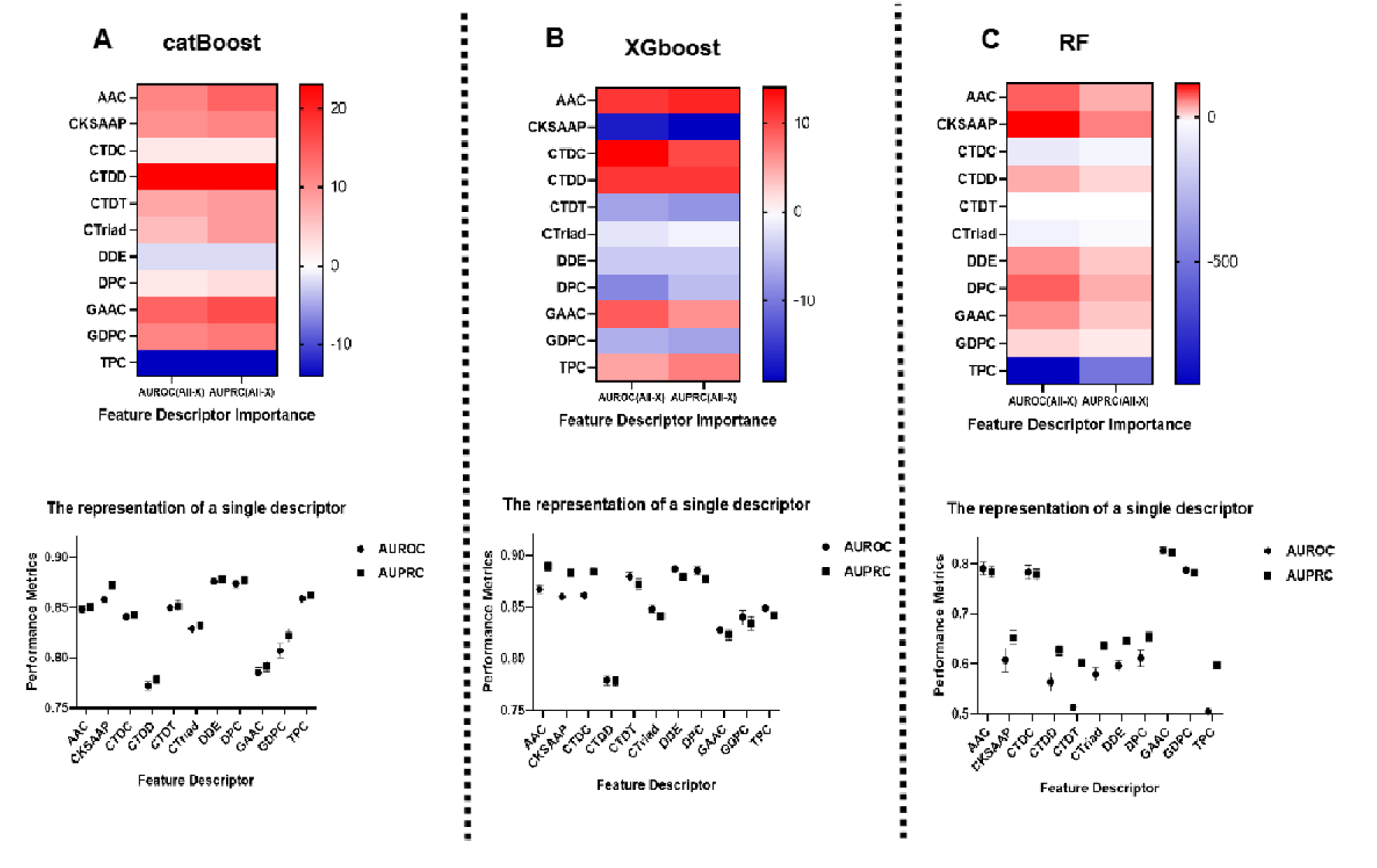
Importance and performance comparison of each descriptor for different machine learning model on 5-Fold Cross-Validation. (A) Importance and performance of feature descriptors based on catBoost; (B) Importance and performance of feature descriptors based on XGBoost; (C) Importance and performance of feature descriptors based on RF.

To assess the contribution of different descriptors to halophilic protein predictions, we initially adopted an iterative approach to systematically exclude individual descriptors from three different models including catBoost(13), XGBoost(14), and Random Forest(15). Since the dataset is imbalanced, the precision is insufficient to correctly evaluate the performance of the model; therefore, AUROC(16) (Area Under the Receiver Operating Characteristic Curve) and AUPRC(17)(Area Under the Precision-Recall Curve) were used as the main evaluation matrix. We then compared the difference in AUROC and AUPRC between models trained with the full descriptor set and models that omitted specific descriptors. Furthermore, we calculated the AUROC value and AUPRC value of each descriptor when trained individually, thereby quantifying the importance of each descriptor.

To illustrate feature importance analysis, we could take the Amino Acid Composition (AAC) descriptor as an example. The calculated formula of its importance score is: AAC Importance = [AUPRC (All Features) – AUPRC (All - AAC)] x 10000, where AUPRC refers to the area under the precision-recall curve. This essentially quantified the extent to which model performance degrades when AAC is removed from the full feature set. A positive importance score indicates that AAC improves the model, while a negative score means that it worsens performance. We applied this methodology to calculate the importance of each descriptor for each machine learning model separately. For a given model, if a descriptor had a positive importance score and an individual AUROC and AUPRC exceeding 0.6, we considered it to have contributed positively to the model. Descriptors that do not meet these criteria were deemed to have a negative contribution.

To evaluate the contribution of various predictors to halophilic protein prediction, we quantified the importance of the descriptors by iteratively eliminating single descriptors from the three models and comparing the difference in AUROC values and AUPRC values. These results indicated that using all descriptors to obtain features might not be the best approach for all models. For catBoost, TPC and DDE, feature descriptors had a negative impact on prediction (**Fig. 2A, Table S1**). For XGBoost, CKSAAP, CTDT, CTriad, DDE, DPC and GDPC, feature descriptors had a negative impact on prediction (**Fig. 2B, Table S2**). For RF, CTDC, CTDT, CTirad, DDE and TPC had a negative impact on the prediction (**Fig. 2C, Table S3**).

Therefore, we eliminated descriptors with negative contributions and subsequently recoded using new descriptors for each model with a 5-fold cross-validation method. The results showed that after removing the respective descriptors, the performance of catBoost model was improved (**Table S1**), and the training speed was significantly increased and the calculation time was reduced. Similarly, the performance of XGBoost and RF has also been improved accordingly. Therefore, we confirmed that the model with the best features was the best model based on the 5-fold cross-validation method.

**Figure 3.**
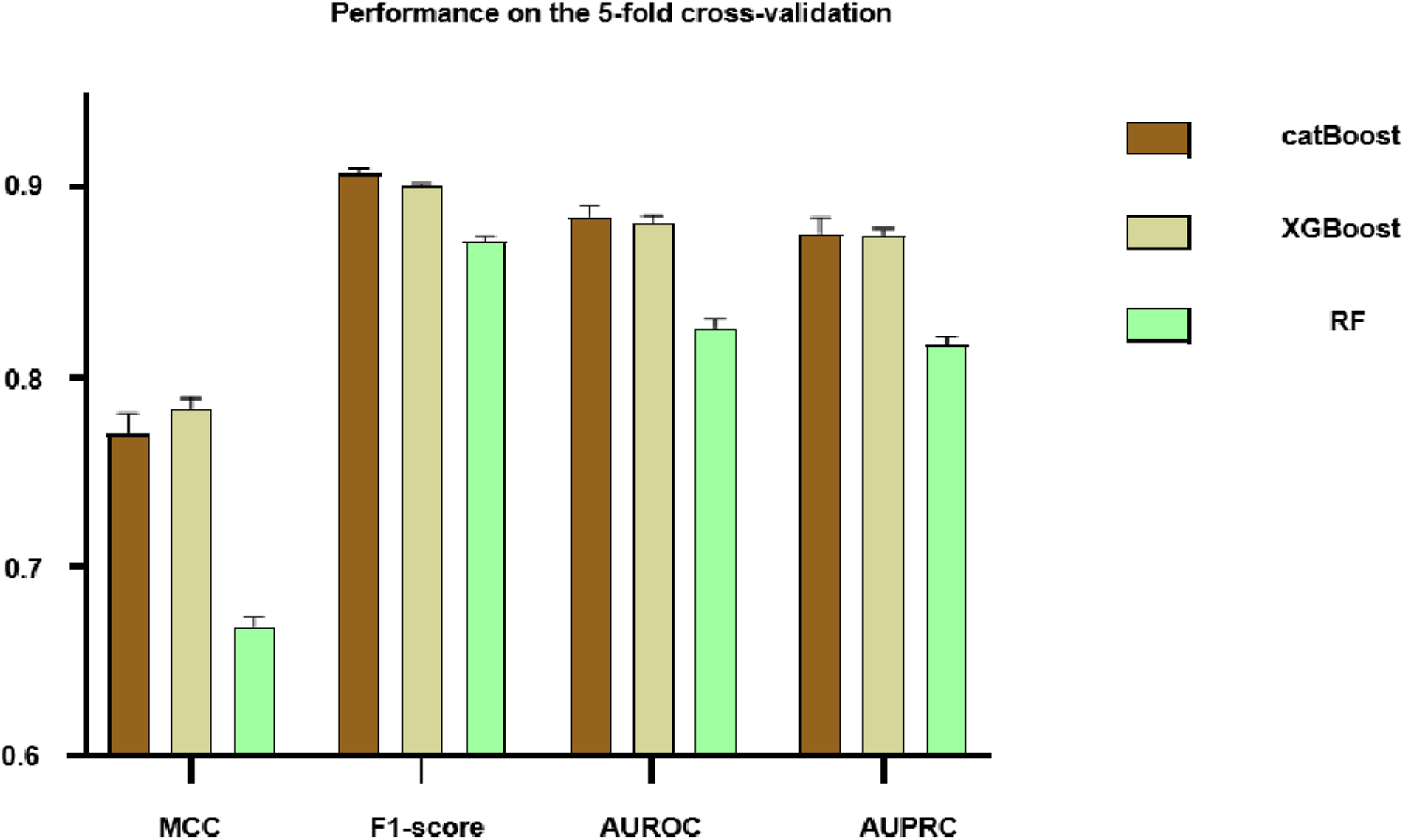
Performance comparison of three models in 5-fold cross-validation after removing descriptors with negative contributions.

Next, we compared the best performance of the three models with the best feature descriptors, as shown in **Figure 3**. Obviously, catBoost outperformed the other models in all evaluation matrices. Moreover, the AUROC and AUPRC of catBoost exceeded 0.8845 and 0.8762 with a very small standard deviation of only 0.0062 and 0.0074 (**Table S1**). This observation further indicated that catBoost not only had high prediction performance but also high robustness. Therefore, catBoost was selected as our main model and a combination of nine descriptors, including AAC, CKSAAP, CTDC, CTDT, CTDD, CTriad, DPC, GAAC and GDPC, was retained for training the final catBoost model.

During the descriptor selection phase of the catBoost training data, we observed that deleting one or two feature descriptors had minimal impact on the predictions of the catBoost model, with AUROC and AUPRC fluctuations remaining below 0.01. This phenomenon could be attributed to three potential reasons: (1) Noise in certain features: Some features might introduce noise and made a small contribution, or even had a negative impact on the prediction performance of the model. Removing these features simplified the model, making it easier to train while maintaining high performance. (2) Robustness of the catBoost model: The catBoost model we adopted shows strong generalization ability and could maintain high performance even when the number of features was reduced(13). (3) Interactions between features: Although some features were less effective when used alone, they might have a significant impact when combined with other features during the training process. Therefore, it might not be sufficient to rely solely on the individual performance of features in the selection process. This could be explained by catBoost’s excellent ability to handle high-dimensional sparse data and feature selection. Removing high-dimensional feature descriptors (such as TPC with 8,000 features) could notably enhanced the performance of the model. This improvement was particularly significant for model performance given that the initial training sample size exceeded 20,000 data points (18).

Based on extensive research on halophilic proteins, it was found that amino acid composition plays an important role in distinguishing halophilic and non-halophilic proteins (2). The results of feature selection in this study further validated this observation. As shown in **Figure 2**, using descriptors such as AAC, CKSAAP, or DPC alone can produce effective predictive outcomes.

It must be acknowledged that the use of handcrafted features primarily based on amino acid composition and some physicochemical properties to train the model may pose several challenges, such as the difficulty in distinguishing site-directed mutations. This could significantly reduce the usefulness of the method in halophilic protein engineering. The focus of study is to collect a rich dataset and train a highly accurate classifier model. In the future, we hoped to employ more comprehensive datasets and better algorithms, such as graph neural networks or pre-trained large language models or even interpretable models, to accomplish the design task of halophilic proteins.

### 2.3 Further Optimization of the Model for Enhanced Performance

In this study, we aimed to enhance the model performance of the catBoost model by employing four feature selection methods, including chi-square (Chi2), L1-based feature selection, tree-based feature selection, and variance threshold feature selection. The optimal feature selection method was determined based on the results obtained from 5-fold cross-validation and independent testing.

Although the Chi2 feature selection method did not yield the highest Matthew correlation coefficient (MCC), F1 score, AUROC, and AUPRC among the four feature selection methods (**Fig. 4A, Table S4**), the model demonstrated a notably strong performance on the independent test set (**Fig. 4B, Table S5**). This outcome indicated that the Chi2 feature selection method had a better generalization ability.

**Figure 4.**
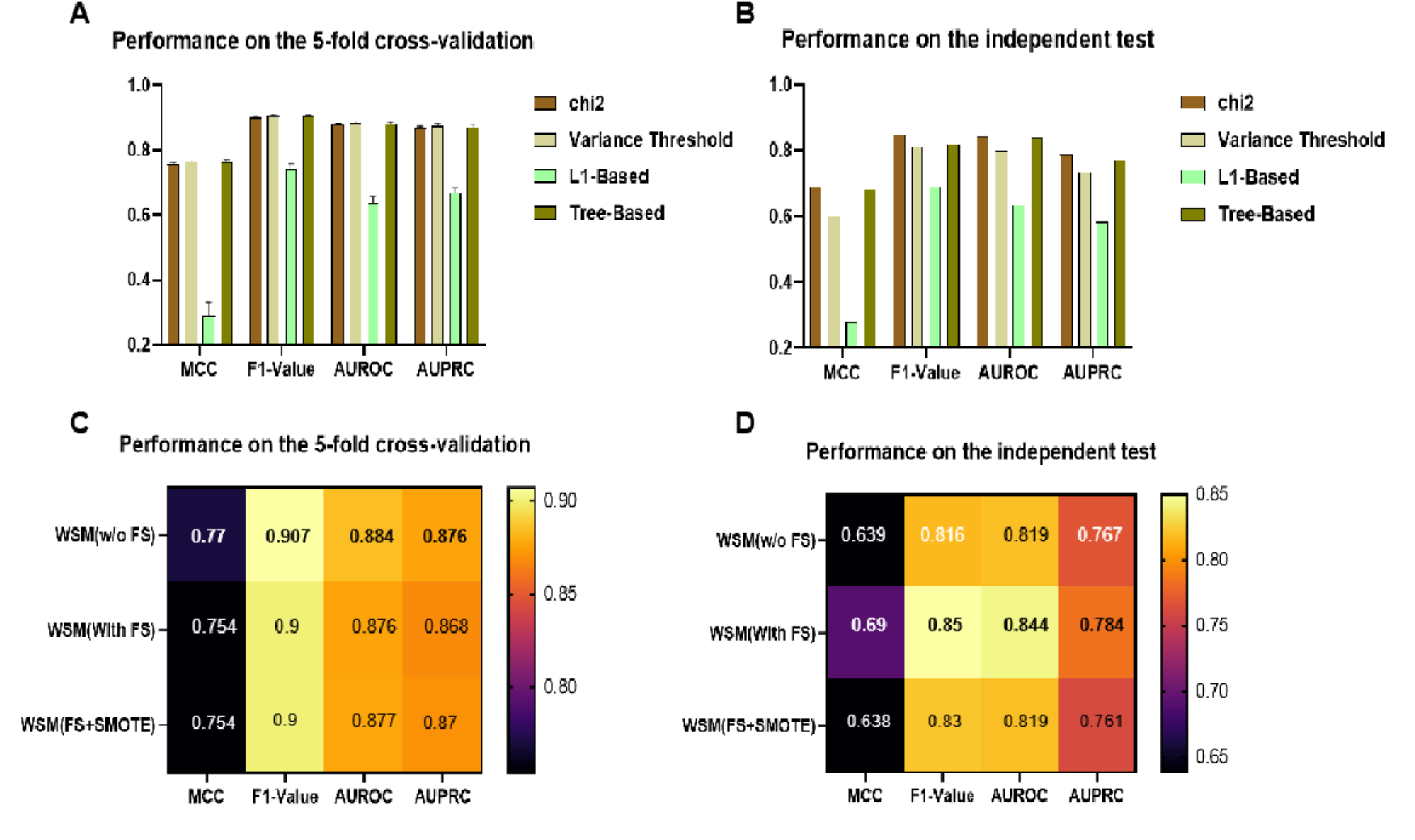
Performance evaluation of different optimization methods. (A) Performance evaluation of different feature selection methods on 5-fold cross-validation; (B) Performance evaluation of different feature selection methods on independent testing; (C) Performance comparison of catBoost without FS, with FS, and with FS + SMOTE on 5-fold cross-validation; (D) Performance comparison of catBoost with different processing methods in independent testing. “WSM” means “whole sequence model” . “ FS” means “ feature selection”. The performance was evaluated based on catBoost model with screened features.

We combined all handcrafted features encoded by eight descriptors with the catBoost model and evaluated the performance of the catBoost model in five-fold cross-validation (**Table S6**) and independent testing (**Table S7**) by integrating the selected features using the Chi2 feature selection strategy. Although the performance of the model dropped slightly after using the feature selection method (**Fig. 4C**), we could see improvements in Matthew correlation coefficient (MCC), F1 score, AUROC, and AUPRC on the independent test set (**Fig. 4D**). This suggested that feature selection indeed improved the generalization ability of the catBoost model.

Considering the data imbalance in the data set, we attempted to solve this issue by using the SMOTE algorithm to balance the data set and applied Chi2 feature selection to identify the best features. However, after introducing SMOTE into the model, it is evident from **Figure 4C** and **Figure 4D** that a significant darker color of the squares was observed compared to when using feature selection alone, indicating the performance of the model was significantly reduced based on 5-fold cross-validation and independent testing. Instead, we used the WSM model with feature selection (denoted WSM(FS) in **Fig. 4C**) as our final Halophilic Protein Classifier (HPClas) model. Despite the class imbalance, this model achieved strong predictive performance without using class balancing techniques such as SMOTE. In summary, feature selection provided a high-quality model, while SMOTE oversampling performed poorly for halophilic protein classifier.

To prove the superiority of the machine learning method we provided, we further conducted a comparative analysis of the HPClas model the commonly used BLASTP and HMM methods on an independent test set. The results are listed in the Table 2. Our machine learning method achieved an accuracy of 84.5% on a test set containing 100 positive and negative samples. In comparison, the accuracy rates of the other two methods were only 64.5% and 64% respectively.

**Table 2.**
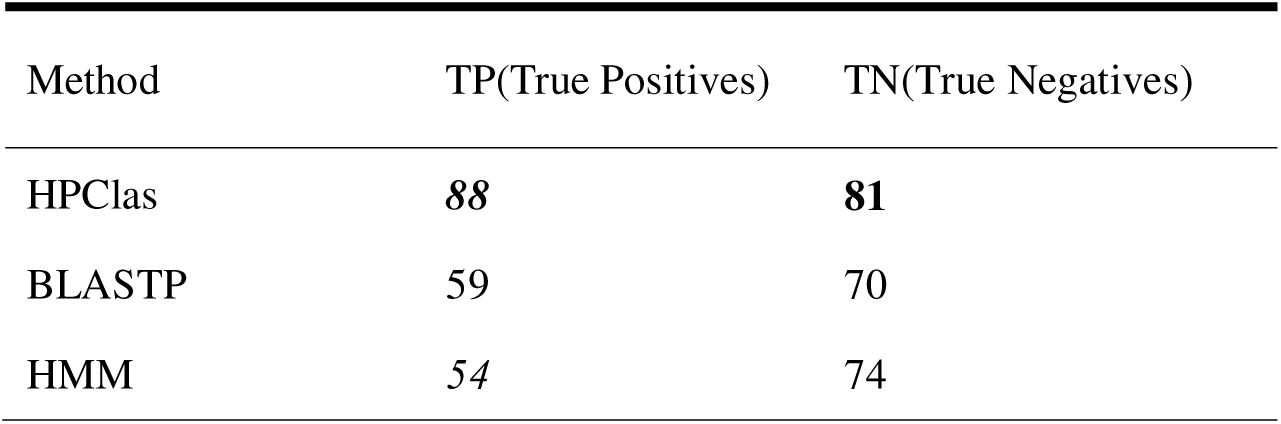
Performance comparison of the three methods on independent test data sets.

| Method | TP(True Positives) | TN(True Negatives) |
| --- | --- | --- |
| HPClas | **88** | **81** |
| BLASTP | 59 | 70 |
| HMM | 54 | 74 |

However, due to the persistent imbalance of more positive samples than negative samples in the dataset, false negatives still exist. Therefore, to meet the needs of different users, we have introduced an adjustable threshold in the prediction setting. Only when the predicted probability is above this threshold will the protein be classified as a halophilic protein. Based on this optimization, users can set thresholds according to their own needs, reduce the false positive rate, and improve the success rate and efficiency of wet laboratory experiments.

### 2.4 Performance comparison with SOTA method on previous test set

To further evaluate the performance of our model, we compared HPClas with other state-of-the-art halophilic protein prediction models on the same data set in **Table 3**. The data set came from the study of Zhang et al. (8, 9), consisting of 15 pairs of halophilic and non-halophilic proteins. Additionally, we have confirmed that these 30 proteins did not appear in our previous training data set. We employed the HPClas model to predict labels for all proteins in the test set and compared the results with those reported in previous works. Experimental results indicated that the HPClas model predicted halophilic proteins with an accuracy of 100% and predicted non-halophilic proteins with an accuracy of 100%. In comparison, previous earlier studies utilizing Artificial Neural Networks (ANN) and Support Vector Machines (SVM) achieved accuracies of 80% and 73%, respectively (8, 9). These findings demonstrated that the HPClas model outperformed previous models in predicting both halophilic and non-halophilic proteins, and exhibiting higher accuracy.

**Table 3.** Performance comparison of the two methods on the previous testing data set.

| PDB ID | Actual type | Predicted type |  |
| --- | --- | --- | --- |
|  |  | HPClas | Previous |
| 1DOI:A | HP | HP | HP |
| 1NWZ:A | HP | HP | NHP |
| 1TJO:A | HP | HP | HP |
| 2B5W:A | HP | HP | HP |
| 2CC6:A | HP | HP | HP |
| 3IBM:A | HP | HP | HP |
| 1ITK:A | HP | HP | HP |
| 2AZ3:A | HP | HP | HP |
| 2J5K:A | HP | HP | HP |
| 1CNO:A | HP | HP | NHP |
| 1NML:A | HP | HP | NHP |
| 2VPN:A | HP | HP | HP |
| 3IFV:A | HP | HP | HP |
| 3IGN:A | HP | HP | HP |
| 3BSM:A | HP | HP | HP |
| 1FXA:A | NHP | NHP | NHP |
| 1MZU:A | NHP | NHP | HP |
| 2VXX:A | NHP | NHP | NHP |
| 2CD9:A | NHP | NHP | NHP |
| 2V18:A | NHP | NHP | NHP |
| 3KGZ:A | NHP | NHP | HP |
| 2FXG:A | NHP | NHP | HP |
| 3B54:A | NHP | NHP | NHP |
| 1Y6J:A | NHP | NHP | NHP |
| 1ETP:A | NHP | NHP | NHP |
| 3HQ6:A | NHP | NHP | NHP |
| 3FXB:A | NHP | NHP | NHP |
| 1RWZ:A | NHP | NHP | NHP |
| 3I5C:A | NHP | NHP | NHP |
| 2QJJ:A | NHP | NHP | HP |

In previous studies, the researchers initially collected positive and negative samples from two bacterial strains, *S. ruber* DSM 13,855 and *P. luteolum* DSM 2379, to form a training set. They used amino acid composition as a feature representation method and used artificial neural networks (ANN) and support vector machine (SVM) to achieve the optimal prediction performance. However, none of them could accurately predict the label of PDB IDs 1NWZ and 1CNO, which were photoreceptor PYP (photoactive yellow protein) (19) and cytochrome c552 (20), respectively. One potential reason was that these proteins were membrane proteins with different amino acid compositions than other proteins. This disparity might lead to inaccurate predictions from previous models. In comparison, our model successfully identified these proteins and accurately predicted them, demonstrating the comprehensiveness of our data sources and the accuracy of the trained model.

To make our work more accessible, we developed the model as a local standalone tool and publish it publicly on GitHub. Additionally, to meet different needs, we incorporated multiple flexible threshold options to the local standalone version of HPClas. Users can adjust these thresholds to prioritize precision or recall in predictions. By sharing the HPClas model code and providing adjustable threshold options, we emphasized the transparency and reproducibility of our approach and aimed to promote collaboration and communication between the academic and bioinformatics communities. We believe that this open and flexible approach should facilitate further advances in the research and discovery of halophilic proteins.

### 2.5 Performance of HPClas Model on Real Halophilic Proteins

To evaluate the practical utility of HPClas in a real-world setting, we tested 16 experimentally validated halophilic enzymes as shown in **Table 4**. We retrieved their amino acid sequences from the RCSB Protein Data Bank (PDB) database based on the accession codes provided. These sequences were used as queries against the trained HPClas model to predict whether they would be classified as halophilic or non-halophilic.

**Table 4.** Performance of HPClas in Experimental Data.

| Halophilic enzymes | Source | PDB number | Optimal and tolerable salt concentration | Predicted type | References |
| --- | --- | --- | --- | --- | --- |
| Cellulase | *Bacillus* sp. BG-CS10 | 5EOC | Optimal 2.5 mol/L NaCl | HP | (21) |
| Carbonic anhydrase | Bovine | 4CNR | Tolerable 3.0 mol/L NaCl | HP | (22) |
| Carbonic anhydrase | *Dunaliella* | 1Y7W | Tolerable 2.0 mol/L NaCl | HP | (23) |
| Carbonic anhydrase | *Photobacterium profundum* | 5HPJ | Optimal 0.5 mol/L NaCl | NHP | (24) |
| Alkaline phosphatase | *Halomonas* sp. 593 | 3WBH | Tolerable 1.0–4.0 | NHP | (25) |

|  |  |  | mol/L NaCl |  |  |
| --- | --- | --- | --- | --- | --- |
| Malate dehydrogenase | *Salinibacter ruber* | 4CL3 |  | HP | (26) |
| Malate dehydrogenase | *Haloarcula marismortui* | 4JCO |  | HP | Not Published |
| Rnase H1 | *Halobacterium salinarum* | 4NYN | Tolerable<br>3.0 mol/L NaCl | HP | Not Published |
| Nucleoside diphosphate kinase | *Haloarcula quadrata* | 2ZUA | Tolerable<br>0.2–4.0 mol/L NaCl | HP | (27) |
| Nucleoside diphosphate kinase | *Halomonas sp. 593* | 3VGS | Tolerable<br>2.0 mol/L NaCl | HP | (28) |
| Malate synthase | *Haloferax volcanii* | 3OYX | Tolerable<br>3.0 mol/L KCl | HP | (29) |
| Endonuclease | *Aliivibrio salmonicida* | 2PU3 | Optimal<br>0.4 mol/L NaCl | NHP | (30) |
| α-Amylase | *Halothermothrix orenii* | 3BC9 | Optimal<br>0.9 mol/L NaCl | HP | (31) |
| α-Amylase | *Halothermothrix orenii* | 1WZA | Tolerable<br>4.7 mol/L NaCl | HP | (3) |
| DHFR | *Haloferax volcanii* | 2ITH | Tolerable<br>3.5 mol/L NaCl | HP | (32) |

Of the 16 known halophilic proteins, HPClas correctly predicted 13 of them as halophilic and 3 of them as non-halophilic. The results demonstrated that HPClas could effectively (albeit imperfectly) identify true halophilic proteins in many cases. There were still some false negative predictions where known halophilic proteins were misclassified. Nonetheless, predictions of true halophilic proteins proved that HPClas has learned relevant patterns and could generalize beyond the training data to identify halophilic proteins in practice. Future testing on a more diverse data set of halophilic proteins could further validate the practical applicability of this model.

### 2.6 Feature Importance Analysis

Importance analysis using the SHAP method allowed us to better understand the decision-making process of the model and the contributions of each feature to the prediction results (33). This enabled us to make more targeted adjustments in feature selection and model optimization, improving the performance and robustness of the model. In addition, importance analysis could help us identify potential issues, such as redundant features and interrelationships between features, which need to be focused on during model development (34).

**Figure 5.**
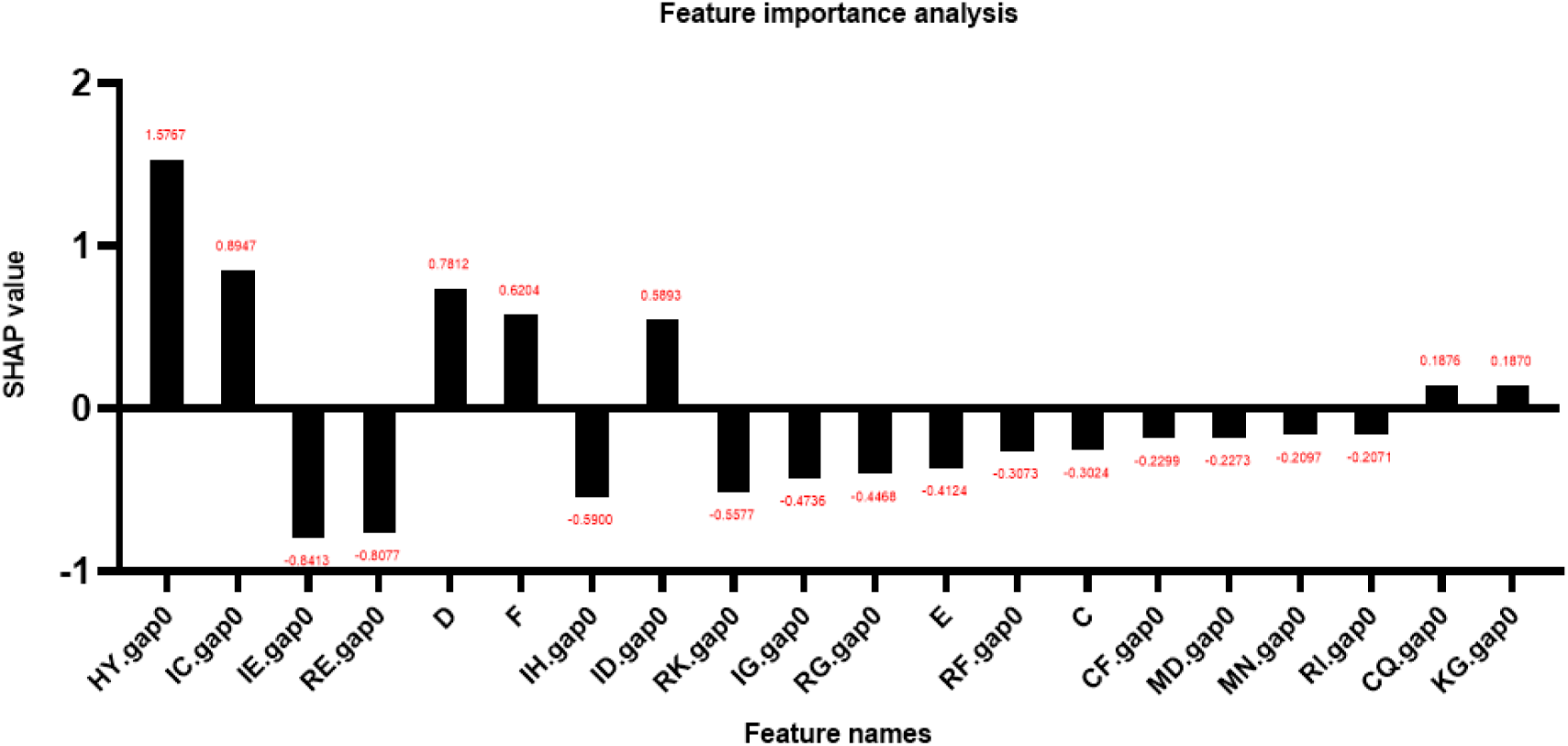
Feature importance analysis. It displays the statistical scores for each feature of the input obtained through SHAP analysis. “XY.gap0”, where X and Y represent two different amino acids, typically denotes a count of zero-gap amino acid pairs that shows variability within a protein sequence. In “X”, X represents a specific amino acid, and this feature indicates the frequency of occurrence of the amino acid represented by X in the entire protein.

In addition, this study also conducted a coarse-grained analysis of the model’s prediction process. We evaluated the importance of each handcrafted input feature for classification and presented partial results in **Figure 5**. Through the charts, we could clearly observe the crucial role of dipeptide combinations and the frequency of certain amino acids in classification decisions. Regarding the single amino acid content, it is evident that the presence of aspartic acid significantly affected positive prediction, while glutamic acid had a substantial impact on negative prediction. These findings were consistent with a 2014 statistical study of halophilic proteins by Giuseppe Graziano and colleagues.

As for dipeptide-type features, it is noteworthy that dipeptide features dominated the top ranking positions. Over the past three decades, protein dipeptide composition has played an important role in predicting ion channels(35), protein structural categories (36), subcellular localization (37), nuclear receptors (38), protein thermal stability (39), non-classical protein secretion (16), protein secondary structure content (40), and more. In this study, although a direct presentation of dipeptide feature descriptors was not provided, the identification of dipeptide features within the CKSAAP descriptor through feature selection indirectly highlighted the importance of these features in the field of halophilic protein prediction.

The interpretability analysis results of this study were consistent with the reported properties of halophilic proteins. Although we could not directly deduce all the characteristics of halophilic proteins and directly explain the identification process based on biological features, the results obtained were sufficient to prove that during the prediction process, the model did not rely on the “memory” training set, but truly learn halophilic-related features and make predictions based on these features. This further confirmed the reliability of the model.

Through fine-grained interpretability analysis of the model’s prediction process, we gained insights into how the model made predictions and the features on which its decision depended. This not only enhanced our understanding of the model prediction results but also provided clues and inspiration for further biological research. We believed that this study will make valuable contributions to the identification and application of halophilic proteins.

## 3. Materials and methods

### 3.1 Collection of Halophilic Protein Data

#### 3.1.1 Collection of Training Sets

To obtain halophilic protein sequences, we searched the NCBI protein database (41) using the keywords “halophilic bacteria” and “halophilic archaea”. Subsequently, SignalP 6.0 software was used to predict signal peptides for the resulting proteins (42), and those lacking signal peptides were excluded. Then, The TMHMM(43) online tool was used to predict the transmembrane regions, thereby excluding proteins containing transmembrane domains. The remaining protein, which contains a signal peptide and lacks a transmembrane region, was identified as a halophilic protein. Similarly, proteins with signal peptides from freshwater microorganisms were considered as non-halophilic proteins. The collected redundant proteins were removed using CD-HIT(44) with a sequence identity threshold of 40%. Also, the protein length was restricted to between 150 and 4000. These data collection and processing steps resulted in two diverse and comprehensive data sets, including halophilic and non-halophilic proteins.

#### 3.1.2 Collection of Independent Test Sets

In terms of the test data set, we primarily used the keywords “Halophilic bacteria” to search for halophilic proteins from UniProtKB and collected them as halophilic proteins. Then, the protein sequences of *Chryseobacterium* NBC 122 were collected from UniProtKB (45) as a non-halophilic sample because NBC 122 isolated from fresh water exhibited optimal growth conditions in 1% NaCl and stops growing above 3.5% NaCl (46). All sequences were pretreated by several steps, including redundancy reduction, length restriction, signal peptide prediction, and transmembrane region prediction. MMseqs2(47) were used concurrently to ensure the homology of samples in the test and training sets remained below 25%.

### 3.2 Feature engineering

#### 3.2.1 Feature Descriptors

The employed 11 feature descriptors include: (1) AAC (Amino Acid Composition): This descriptor counts the frequency of each amino acid in a protein sequence; (2) CKSAAP (Composition of K-spaced Amino Acid Pairs): It counts the frequency of amino acid pairs with a specific spacing (K) between them; (3) DPC (Dipeptide Composition): This descriptor examines the frequency of consecutive dipeptides in a protein sequence; (4) TPC (Tri-peptide Composition): It counts the frequency of amino acid tripeptides in the sequence; (5) CTDC (Composition): This descriptor characterizes the composition of a protein sequence based on three physicochemical groups; (6) CTDT (Transition): The CTDT descriptor measures the percentage frequency of amino acids following a specific amino acid belonging to a different physicochemical group; (7) CTDD (Distribution): It describes the distribution of each physicochemical group along the protein sequence; (8) CTriad (Correlation of Triplet Amino Acids): This descriptor maps protein sequence into a frequency matrix based on amino acid triplets; (9) GAAC (Grouped Amino Acid Composition): It calculates the frequency of amino acids based on predefined amino acid groups; (10) GDPC (Grouped Dipeptide Composition): This descriptor calculates the frequency of dipeptides based on predefined amino acid groups; (11) DDE (Dipeptide Deviation from Expected): It examines the deviation between the observed frequency of dipeptides and the expected values in the protein sequence. All feature descriptors used here could be calculated using feature evolution and machine learning tools such as iFeature (48), iLearn (49), and iLearnPlus (50). A detailed description of each feature descriptor is provided in the supplementary material.

#### 3.2.2 Feature normalization

For normalizing the values of different features, MinMaxScaler was used from the scikit-learn package (51). The normalization function could be formulated as follows:

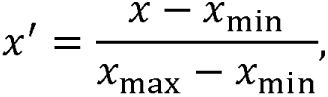

where x, x_mIn_, x_max_ denote the original value, minimum value and maximum value in the feature vectors, respectively, and x^i^ denotes the scaled feature.

#### 3.2.3 Feature selection

Several feature selection methods were employed to construct the model, including chi-square (Chi2) (52), L1-based feature selection (53), tree-based feature selection (54), and variance threshold feature selection (55). Chi2 feature selection is a univariate method that selects the best k features based on the chi-square statistic (52). L1-based feature uses a linear model and L1 normalization penalty to eliminates feature with zero coefficients (53). Tree-based feature selection calculates importance based on impurities and discards irrelevant features (54). Variance Threshold is a simple procedure based on feature variance that discards features below a certain variance threshold (55). All four feature selection methods could be implemented using the “feature_selection” module in scikit-learn.

### 3.3 Model Construction and Optimization

#### 3.3.1 Model Construction

catBoost and XGBoost represent gradient boosting tree models, while Random Forest is a tree-based ensemble model(15). During the training process of the first two models, performance enhancement was achieved by iteratively training a series of decision trees. In contrast, the Random Forest model constructed multiple independently trained decision trees and made predictions through collective voting(56).

When building these models, Python libraries associated with these three models were employed. Initially, a classifier pipeline was built, including data preprocessing steps such as feature scaling (MinMaxScaler), feature selection (SelectKBest), and oversampling (SMOTE), as well as the classifiers themselves. In the classifier section, various parameters of each classifier were configured, including the number of iterations, learning rate, tree depth, and regularization parameters. Subsequently, the models were fitted using the training data, resulting in a trained classifier. Detailed model information is documented in the supplementary materials, and the code and specific hyperparameter values used for model training are publicly available in trainModel.py at https://github.com/Showmake2/HPClas.

#### 3.3.2 The Synthetic Minority Oversampling Technique (SMOTE)

Synthetic Minority Oversampling Technique (SMOTE) (57) was used to treat the class imbalance in our data sets. SMOTE operates by selecting samples from the minority class, identifying their nearest neighbors, and generating synthetic similar samples to increase the number of minority cases. This oversampling is implemented in the “imblearn” Python package.

#### 3.3.3 Randomized 5-Fold Cross-Validation

The benchmark data set was randomly divided into five equal subsets. The evaluation process was repeated five times, with each subset used once as testing data and four times for training. In each iteration of cross-validation, one subset was reserved for testing, while the remaining four subsets were combined into a training set for training the classifier. The results of the five tests were then averaged to obtain a single performance value that represents the overall performance of the classification model. Additionally, hyperparameter tuning was conducted using a grid search approach (58). Scikit-learn’s KFold method (59) was then used to implement 5-fold cross-validation, and the performance metrics of the model were present as mean ± standard deviation.

#### 3.3.4 Performance assessment

The seven performance indicators commonly used to evaluate model performance include precision, recall, accuracy, Matthew correlation coefficient (MCC) (60), F1 score, area under the receiver operating characteristic curve (ROC) (AUROC), and area under the precision-recall curve (PR) (AUPRC). These performance metrics can be calculated as follows:

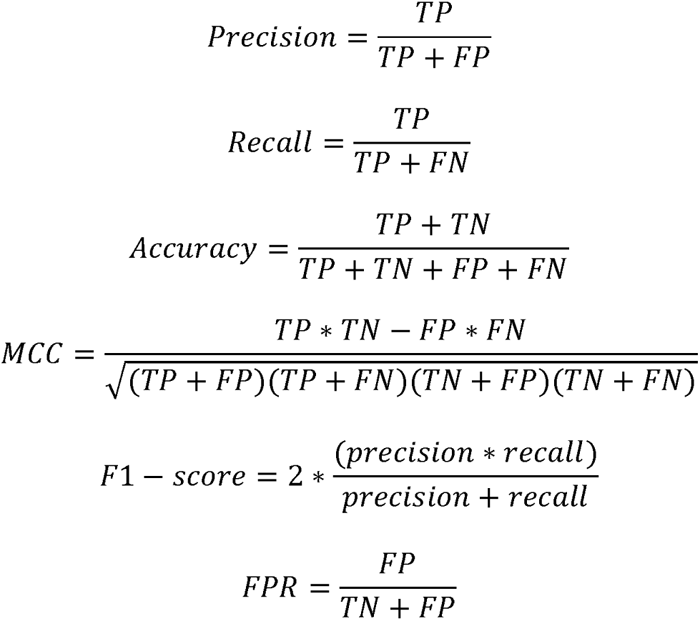

where TN, TP, FN, and FP represent the number of true negatives (i.e., correctly predicted non-halophilic proteins), true positives (i.e., correctly predicted halophilic proteins), false negatives (i.e., incorrectly predicted non-halophilic proteins), and false positives (i.e., incorrectly predicted halophilic proteins), respectively. The x-axis of the ROC curve represents the false positive rate (FPR) and the y-axis represents the true-positive rate (TPR). The precision-recall curve plots precision on the y-axis and recall on the x-axis. Consequently, the area under the precision-recall curve was also used as the primary metric to evaluate the performance of different models when the data set is highly imbalanced. Since the dataset is imbalanced, the precision is insufficient to correctly evaluate the performance of the model; therefore, AUROC and AUPRC were used as the main evaluation matrix.

### 3.4 Feature Importance Analysis

The SHAP (SHapley Additive exPlanations) value method was employed to evaluate the importance score of each input feature predicted by the model (33). Specifically, for a data point with N features, there exist N! possible ordering or permutations of those features. SHAP evaluates the marginal contribution of each feature by examining how the model output changes when a feature is added to an existing feature set. This is done for all possible feature permutations to account for interaction effects between features. The marginal contribution of a feature in each permutation was weighted by the probability of that permutation occurring. Finally, the SHAP value for each feature was computed as the weighted average of the marginal contributions of all permutations. In essence, the SHAP values explain how much each feature contributes to the model output (either positively or negatively). This provides interpretability of how the model depends on specific input features.

## Acknowledgments

This work was supported by National Key R&D Program of China (NO.2023YFC3403600), National Natural Science Foundation of China (32370101) and Fundamental Research Funds for the Central Universities (buctrc202131).

## Ethics statement

The study in this article did not involve any trials on humans or animals.

## Code and Data Availability

The code and dataset can be accessed at https://github.com/Showmake2/HPClas.

## References

1. Ling C, Qiao GQ, Shuai BW, Olavarria K, Yin J, Xiang RJ, et al. Engineering NADH/NAD(+) ratio in Halomonas bluephagenesis for enhanced production of polyhydroxyalkanoates (PHA). Metab Eng. 2018;49:275–86.

2. Graziano G, Merlino A. Molecular bases of protein halotolerance. Biochim Biophys Acta. 2014;1844(4):850–8.

3. Sivakumar N, Li N, Tang JW, Patel BK, Swaminathan K. Crystal structure of AmyA lacks acidic surface and provide insights into protein stability at poly-extreme condition. FEBS Lett. 2006;580(11):2646–52.

4. Sinha R, Khare SK. Effect of organic solvents on the structure and activity of moderately halophilic Bacillus sp. EMB9 protease. Extremophiles. 2014;18(6):1057–66.

5. Littlechild JA. Enzymes from Extreme Environments and Their Industrial Applications. Front Bioeng Biotechnol. 2015;3:161.

6. Sharma N, Farooqi MS, Chaturvedi KK, Lal SB, Grover M, Rai A, et al. The Halophile protein database. Database (Oxford). 2014;2014:bau114.

7. Gunde-Cimerman N, Plemenitaš A, Oren A. Strategies of adaptation of microorganisms of the three domains of life to high salt concentrations. FEMS Microbiology Reviews. 2018;42(3):353–75.

8. Zhang G, Ge H. Protein hypersaline adaptation: insight from amino acids with machine learning algorithms. Protein J. 2013;32(4):239–45.

9. Zhang G, Huihua G, Yi L. Stability of halophilic proteins: from dipeptide attributes to discrimination classifier. Int J Biol Macromol. 2013;53:1–6.

10. Boutet E, Lieberherr D, Tognolli M, Schneider M, Bansal P, Bridge AJ, et al. UniProtKB/Swiss-Prot, the manually annotated section of the UniProt KnowledgeBase: how to use the entry view. Plant bioinformatics: methods and protocols. 2016:23–54.

11. Nielsen H. Predicting Secretory Proteins with SignalP. Methods Mol Biol. 2017;1611:59–73.

12. Makhdoumi-Kakhki A, Amoozegar MA, Ventosa A. Salinibacter iranicus sp. nov. and Salinibacter luteus sp. nov., isolated from a salt lake, and emended descriptions of the genus Salinibacter and of Salinibacter ruber. Int J Syst Evol Microbiol. 2012;62(Pt 7):1521–7.

13. Prokhorenkova L, Gusev G, Vorobev A, Dorogush AV, Gulin A. CatBoost: unbiased boosting with categorical features. NIPS. 2018;31.

14. Ogunleye A, Wang Q-G. XGBoost model for chronic kidney disease diagnosis. IEEE ACM T COMPUT BI 2019;17(6):2131–40.

15. Breiman L. Random forests. Mach Learn. 2001;45:5–32.

16. Wang X, Li F, Xu J, Rong J, Webb GI, Ge Z, et al. ASPIRER: a new computational approach for identifying non-classical secreted proteins based on deep learning. Brief Bioinform. 2022;23(2).

17. Wen P, Xu Q, Yang Z, He Y, Huang Q. Exploring the Algorithm-Dependent Generalization of AUPRC Optimization with List Stability. NIPS. 2022;35:28335–49.

18. Bhasin M, Raghava GP. ESLpred: SVM-based method for subcellular localization of eukaryotic proteins using dipeptide composition and PSI-BLAST. Nucleic Acids Res. 2004;32(Web Server issue):W414–9.

19. Imamoto Y, Kataoka M. Structure and photoreaction of photoactive yellow protein, a structural prototype of the PAS domain superfamily. Photochem Photobiol. 2007;83(1):40–9.

20. Brown K, Nurizzo D, Besson S, Shepard W, Moura J, Moura I, et al. MAD structure of Pseudomonas nautica dimeric cytochrome c552 mimicks the c4 Dihemic cytochrome domain association. J Mol Biol. 1999;289(4):1017–28.

21. Sandomenico A, Leonardi A, Berisio R, Sanguigno L, Focà G, Focà A, et al. Generation and Characterization of Monoclonal Antibodies against a Cyclic Variant of Hepatitis C Virus E2 Epitope 412-422. J Virol. 2016;90(7):3745–59.

22. Warden AC, Williams M, Peat TS, Seabrook SA, Newman J, Dojchinov G, et al. Rational engineering of a mesohalophilic carbonic anhydrase to an extreme halotolerant biocatalyst. Nat. Commun. 2015;6(1):10278.

23. Premkumar L, Greenblatt HM, Bageshwar UK, Savchenko T, Gokhman I, Sussman JL, et al. Three-dimensional structure of a halotolerant algal carbonic anhydrase predicts halotolerance of a mammalian homolog. Proc Natl Acad Sci U S A. 2005;102(21):7493–8.

24. Somalinga V, Buhrman G, Arun A, Rose RB, Grunden AM. A High-Resolution Crystal Structure of a Psychrohalophilic α-Carbonic Anhydrase from Photobacterium profundum Reveals a Unique Dimer Interface. PLoS One. 2016;11(12):e0168022.

25. Arai S, Yonezawa Y, Ishibashi M, Matsumoto F, Adachi M, Tamada T, et al. Structural characteristics of alkaline phosphatase from the moderately halophilic bacterium Halomonas sp. 593. Acta Crystallogr D Biol Crystallogr. 2014;70(Pt 3):811–20.

26. Talon R, Coquelle N, Madern D, Girard E. An experimental point of view on hydration/solvation in halophilic proteins. Front Microbiol. 2014;5:66.

27. Yamamura A, Ichimura T, Kamekura M, Mizuki T, Usami R, Makino T, et al. Molecular mechanism of distinct salt-dependent enzyme activity of two halophilic nucleoside diphosphate kinases. Biophys J. 2009;96(11):4692–700.

28. Arai S, Yonezawa Y, Okazaki N, Matsumoto F, Tamada T, Tokunaga H, et al. A structural mechanism for dimeric to tetrameric oligomer conversion in Halomonas sp. nucleoside diphosphate kinase. Protein Sci. 2012;21(4):498–510.

29. Bracken CD, Neighbor AM, Lamlenn KK, Thomas GC, Schubert HL, Whitby FG, et al. Crystal structures of a halophilic archaeal malate synthase from Haloferax volcanii and comparisons with isoforms A and G. BMC Struct Biol. 2011;11:23.

30. Altermark B, Helland R, Moe E, Willassen NP, Smalås AO. Structural adaptation of endonuclease I from the cold-adapted and halophilic bacterium Vibrio salmonicida. Acta Crystallogr D Biol Crystallogr. 2008;64(Pt 4):368–76.

31. Tan TC, Mijts BN, Swaminathan K, Patel BK, Divne C. Crystal structure of the polyextremophilic alpha-amylase AmyB from Halothermothrix orenii: details of a productive enzyme-substrate complex and an N domain with a role in binding raw starch. J Mol Biol. 2008;378(4):852–70.

32. Binbuga B, Boroujerdi AF, Young JK. Structure in an extreme environment: NMR at high salt. Protein Sci. 2007;16(8):1783–7.

33. Mangalathu S, Hwang S-H, Jeon J-S. Failure mode and effects analysis of RC members based on machine-learning-based SHapley Additive exPlanations (SHAP) approach. Eng. Struct.. 2020;219:110927.

34. Adadi A, Berrada M. Peeking inside the black-box: a survey on explainable artificial intelligence (XAI). IEEE access. 2018;6:52138–60.

35. Lin H, Ding H. Predicting ion channels and their types by the dipeptide mode of pseudo amino acid composition. J. Theor. Biol. 2011;269(1):64–9.

36. Lin H, Li Q-Z. Using pseudo amino acid composition to predict protein structural class: Approached by incorporating 400 dipeptide components. J Comput Chem. 2007;28(9):1463–6.

37. Li LQ, Zhang Y, Zou LY, Zhou Y, Zheng XQ. Prediction of protein subcellular multi-localization based on the general form of Chou’s pseudo amino acid composition. Protein Pept Lett. 2012;19(4):375–87.

38. Bhasin M, Raghava GP. Classification of nuclear receptors based on amino acid composition and dipeptide composition. J Biol Chem. 2004;279(22):23262–6.

39. Feng C, Ma Z, Yang D, Li X, Zhang J, Li Y. A Method for Prediction of Thermophilic Protein Based on Reduced Amino Acids and Mixed Features. Front Bioeng Biotechnol. 2020;8:285.

40. Chou K-C. Using Pair-Coupled Amino Acid Composition to Predict Protein Secondary Structure Content. Protein J. 1999;18(4):473–80.

41. Sayers EW, Beck J, Bolton EE, Bourexis D, Brister JR, Canese K, et al. Database resources of the National Center for Biotechnology Information. Nucleic Acids Res. 2021;49(D1):D10–d7.

42. Teufel F, Almagro Armenteros JJ, Johansen AR, Gíslason MH, Pihl SI, Tsirigos KD, et al. SignalP 6.0 predicts all five types of signal peptides using protein language models. Nat Biotechnol. 2022;40(7):1023–5.

43. Krogh A, Larsson B, von Heijne G, Sonnhammer EL. Predicting transmembrane protein topology with a hidden Markov model: application to complete genomes. J Mol Biol. 2001;305(3):567–80.

44. Fu L, Niu B, Zhu Z, Wu S, Li W. CD-HIT: accelerated for clustering the next-generation sequencing data. Bioinformatics. 2012;28(23):3150–2.

45. Boutet E, Lieberherr D, Tognolli M, Schneider M, Bairoch A. UniProtKB/Swiss-Prot: the manually annotated section of the UniProt KnowledgeBase. Plant bioinformatics: methods and protocols: Springer; 2007. p. 89–112.

46. Kim H, Yu SM. Chryseobacterium salivictor sp. nov., a plant-growth-promoting bacterium isolated from freshwater. Antonie Van Leeuwenhoek. 2020;113(7):989–95.

47. Steinegger M, Söding J. MMseqs2 enables sensitive protein sequence searching for the analysis of massive data sets. Nat. Biotechnol. 2017;35(11):1026–8.

48. Chen Z, Zhao P, Li F, Leier A, Marquez-Lago TT, Wang Y, et al. iFeature: a Python package and web server for features extraction and selection from protein and peptide sequences. Bioinformatics. 2018;34(14):2499–502.

49. Chen Z, Zhao P, Li F, Marquez-Lago TT, Leier A, Revote J, et al. iLearn: an integrated platform and meta-learner for feature engineering, machine-learning analysis and modeling of DNA, RNA and protein sequence data. Brief Bioinform. 2020;21(3):1047–57.

50. Chen Z, Zhao P, Li C, Li F, Xiang D, Chen YZ, et al. iLearnPlus: a comprehensive and automated machine-learning platform for nucleic acid and protein sequence analysis, prediction and visualization. Nucleic Acids Res. 2021;49(10):e60.

51. Aksoy S, Haralick RM. Feature normalization and likelihood-based similarity measures for image retrieval. Pattern Recogn Lett. 2001;22(5):563–82.

52. Liu H, Setiono R. Proceedings of 7th IEEE International Conference on Tools with Artificial Intelligence. 1995.

53. Ng AY. Feature selection, L1 vs. L2 regularization, and rotational invariance. Proceedings of the twenty-first international conference on Machine learning; Banff, Alberta, Canada: Association for Computing Machinery; 2004. p. 78.

54. Liu Z, Song J. Comparison of Tree-based Feature Selection Algorithms on Biological Omics Dataset. Proceedings of the 5th International Conference on Advances in Artificial Intelligence; Virtual Event, United Kingdom: Association for Computing Machinery; 2022. p. 165–9.

55. Powell A, Bates D, Van Wyk C, de Abreu D, editors. A cross-comparison of feature selection algorithms on multiple cyber security data-sets. FAIR; 2019.

56. Liaw A, Wiener M. Classification and regression by randomForest. R news. 2002;2(3):18–22.

57. Chawla NV, Bowyer KW, Hall LO, Kegelmeyer WP. SMOTE: synthetic minority over-sampling technique. J Artif Intell Res. 2002;16:321–57.

58. Fuadah YN, Pramudito MA, Lim KM. An Optimal Approach for Heart Sound Classification Using Grid Search in Hyperparameter Optimization of Machine Learning. Bioengineering (Basel). 2022;10(1).

59. Sejuti ZA, Islam MS. A hybrid CNN-KNN approach for identification of COVID-19 with 5-fold cross validation. Sens Int. 2023;4:100229.

60. Matthews BW. Comparison of the predicted and observed secondary structure of T4 phage lysozyme. Biochim Biophys Acta Protein Struct. 1975;405(2):442–51.

