## Supplementary material for "HPClas: A data-driven approach for identifying halophilic proteins based on catBoost": Table S1-S7

### Supplemental Methods

#### Feature engineering

###### Amino acid composition：

The amino acid composition characteristics reflect amino acid types and pairing frequencies.

(1) Amino Acid Composition (AAC)

The AAC encoding counts the frequency of 20 amino acids in a peptide sequence (1). It can show the basic composition of amino acid and has been widely used in the field of bioinformatics (2-4). AAC is defined as follows:

$$f\left( x \right)=\frac{N\left( x \right)}{N}, x\in\{A, C, D,\ldots,Y\}$$

where $N\left( x \right)$ is the number of type *x* amino acid residues, and *N* represents the length of the amino acid sequence.

(2) Composition of k-spaced amino acid pairs (CKSAAP)

The CKSAAP encoding scheme was proposed by Chen in 2007 (5), and has been widely used in many bioinformatics prediction tasks, such as functional protein identification (6, 7) and protein functional site prediction (8, 9). It calculates the frequency of amino acid pairs spaced by *k* (*k*=0, 1, 2, 3 in this study) residues (1, 10), and the output of this descriptor is a 1600-dimensional vector. For example, when *k* = 3, the descriptor is calculated as follows:

$${(\frac{N_{AXXXA}}{N_{total}}, \frac{N_{AXXXC}}{N_{total}}, \frac{N_{AXXXD}}{N_{total}},\ldots, \frac{N_{YXXXY}}{N_{total}})}_{400}, X\in\{A, C, D,\ldots,Y\}$$

where $N_{total}$ is the total number of all amino acid pairs spaced by three residues, $N_{AXXXA}$ refers to the number of AA pairs separated by three residues, and X can be any kind of residue.

(3) Dipeptide Deviation from Expected Mean (DDE)

The Dipeptide Deviation from Expected Mean (DDE) was originally proposed by Saravanan et al. (11), which was computed by three parameters: dipeptide composition ($D_{c}$), theoretical mean ($T_{m}$), and theoretical variance ($T_{v}$) (1). The DDE feature descriptor can be calculated as follows:

$$f\left( r,s \right)=\frac{D_{c}\left( r,s \right)-T_{m}(r,s)}{\sqrt{T_{v}(r,s)}}, r,s\in\left\{ A, C, D,\ldots,Y \right\}$$

$D_{c}\left( r,s \right)$ is the dipeptide composition for the $\left( r,s \right)$ dipeptide, and is defined as follows:

$$D_{c}\left( r,s \right)=\frac{N_{rs}}{N-1}, x,y\in\{A, C, D,\ldots,Y\}$$

where *N* is the length of the sequence and $N_{rs}$ is the number of the dipeptide *rs*. *r* and *s* denote the amino acid types. $T_{m}(r,s)$ is given as:

$$T_{m}\left( r,s \right)=\frac{C_{r}}{C_{N}} \times\frac{C_{s}}{C_{N}}$$

where the $C_{r}$ and $C_{s}$ are the number of the codons corresponding to the amino acid *r* and amino acid *s*. For example, methionine (M) has only one codon, so $C_{M}$ is 1. $C_{N}$ is the total number of codons for all 20 amino acid types (i.e. 61). Moreover, $T_{v}$ is the theoretical mean, defined as:

$$T_{v}\left( r,s \right)=\frac{T_{m}(r,s)(1-T_{m}\left( r,s \right))}{N-1}$$

(4) Di-Peptide Composition (DPC)

Di-peptide composition calculates the frequency of di-peptides in a protein sequence and generates a 400-dimensional vector. It can be defined as follows:

$$f\left( x,y \right)=\frac{N_{xy}}{N-1}, x,y\in\{A, C, D,\ldots,Y\}$$

where *N* is the number of the length of the protein or peptide, and $N_{xy}$ is the number of the *xy* dipeptides.

(5) Tri-Peptide composition (TPC)

TPC encoding has been previously used in several studies (12), which generated fixed feature vectors of 8000-dimensions. TPC is defined as follows:

$$f\left( x,y,z \right)=\frac{N_{xyz}}{N-2}, x,y,z\in\left\{ A, C, D,\ldots,Y \right\}$$

where the *x*, *y*, and *z* are the amino acid types, and $N_{xyz}$ is the number of the tripeptide *xyz*.

###### Physicochemical properties：

(6) Composition (CTDC)

The CTDC descriptor characterizes the composition of amino acid sequences from three physicochemical groups (13) , and is defined as follows:

$$f\left( r \right)=\frac{N(r)}{N}, r\in\left\{ polar,neutral,hydrophobic \right\}$$

where $N\left( r \right)$ is the number of particular amino acid residue type *r* and *r* belongs to a physicochemical group, and *N* is the number of amino acids (i.e., the length of the sequence).

(7) Transition (CTDT)

The CTDT descriptor measures the percentage frequency of a particular amino acid followed by another physicochemical group of amino acids (13). The CTDT descriptor is calculated as:

$$f\left( r,s \right)=\frac{N\left( r,s \right)+N\left( s,r \right)}{N-1},$$

$$r,s\in\left\{ \left( polar,neutral \right),\left( neutral,hydrophobic \right),(hydrophobic,polar) \right\}$$

where $N\left( r,s \right)$ denotes the number of dipeptides that contains two amino acids *r*, *s*, and the *r* and *s* belong to different groups.

(8) Distribution (CTDD)

The CTDD descriptor describes the distribution of each physicochemical property in protein sequence (13). It contains five values for each physicochemical group, corresponding to the position fraction of the entire sequence where the first, 25%, 50%, 75% and 100% residues of a certain group are located.

(9) Conjoint Triad (CTriad)

The CTriad descriptor represents the properties of amino acids and their vicinal amino acids, and three continuous amino acids can be identified as one unit(14). Amino acids are classified into seven categories (g1: AGV, g2: ILFP, g3: YMTS, g4: HNQW, g5: RK, g6: DE, g7: C). It is a 343-dimensional vector, and each vector is the frequency of conjoint triad occurring in the peptides sequence.

###### Grouped amino acid composition：

(10) Grouped amino acid composition (GAAC)

The GAAC encoding scheme calculates the frequency of amino acid types based on their physicochemical properties and is a variant of the AAC descriptor. Amino acids are classified into five categories: aliphatic (group 1: GAVLMI), aromatic (group 2: FYW), positively charged (group 3: KRH), negatively charged (group 4: DE), and uncharged (group 5: STCPNQ) (15).

$$f\left( c \right)=\frac{N\left( c \right)}{N}, c\in\{c1, c2, c3, c4,c5\}$$

$$N\left( C \right)= \sum N(t), t\in c$$

where $N\left( c \right)$ is the number of amino acids in class $c$, and $t$ is the amino acid type belonging to class $c$. *N* denotes the length of the protein.

(11) Grouped Di-Peptide Composition (GDPC)

GDPC coding is similar to DPC in that both calculate the frequency of amino acid pairs, and amino acids are categorized into five classes (16). GDPC returns a 25-dimensional vector. The criteria for amino acid categories are the same as GAAC. It can be defined as:

$$f\left( r,s \right)=\frac{N\left( r,s \right)}{N}, r,s\in\{c1, c2, c3, c4,c5\}$$

$$N\left( r,s \right)= \sum N(t), t \in r,s$$

where $N\left( r,s \right)$ is the number of amino acid pairs $\left( r,s \right)$ and $t$ is the type of amino acid pairs.

CatBoost（Categorical Boosting）

CatBoost is a robust machine learning framework developed by Yandex and launched in 2017 (17). Its key strength is automatic hyperparameter tuning, which simplifies the training process by adapting learning rates. Additionally, CatBoost uses performance optimizations and overfitting prevention mechanisms to make it effective on large-scale datasets.

CatBoost excels at performance optimization, using techniques like symmetric tree growth strategy and query-based gradient boosting. These optimizations help improve its ability to handle large-scale datasets and deliver competitive training and inference speeds (18). The framework also incorporates mechanisms to prevent overfitting, such as statistical learning processes and random tree rotations, ensuring improved generalization of new data (17).

**XGBoost**（eXtreme Gradient Boosting）

XGBoost is a popular and efficient machine learning algorithm library that was born in February 2014, focusing on gradient boosting algorithms. XGBoost is particularly suitable for classification and regression tasks with large-scale datasets and high-dimensional feature spaces. It is a model based on decision tree ensemble learning method (19).

The main advantages of XGBoost are speed and accuracy. It uses ensemble learning through multiple tree models, with each tree model trained on the residuals of the previous tree model. This residual learning approach can effectively reduce the risk of overfitting while improving the accuracy of the model. In addition, XGBoost achieves efficient training and inference speed through techniques such as distributed computing and GPU acceleration.

Random forest

RF algorithm is a classification algorithm developed by Leo Breiman using an ensemble of classification trees. It has been widely used and implemented as the RF package in R. RF is one of the most powerful algorithms in machine learning. In RF, two key parameters are the number of the trees $M$ and the number of randomly selected features mtry (20).

Here, we selected$M=1000$ and optimized the parameter over the set of integers between 1 and

$\lfloor max\{\sqrt{\text{ featureNum}}$, featureNum$/2\}\rfloor$ to minimize the classification error. Here, featureNum is the total number of features.

#### Feature Selection

Chi-square (Chi2)

The chi-square test is a method for hypothesis testing, designed to determine whether two categorical variables are independent of each other (21). It assesses the deviation between observed and expected values to determine the validity of the null hypothesis. The formula of the chi-square test is as follows, where A represents the observed values, corresponding to the data in the first 2x2 table, and T denotes the expected values, which represents the values in the theoretical 2x2 table. The chi-square statistic (X2) measures the difference between observed and expected values and contains two critical pieces of information: the absolute magnitude of the deviation between observed and expected values (amplified by the square) and the relative magnitude of the difference compared to the expected values.

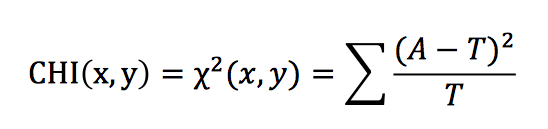

The larger the value of X2, the higher the possibility that the two variables are independent, and the stronger correlation between the two variables. For the feature variables x1, x2, ..., xn, and the categorical variable y, it is necessary to calculate CHI(x1, y), CHI(x2, y), ..., CHI(xn, y), and then the features are sorted based on the CHI values in descending order. Following this, a threshold is chosen such as features with CHI values above the threshold are retained, and features below the threshold are deleted. This process yields a subset of selected features that can then be used to train a classifier, followed by evaluation of the performance of the classifier (22).

L1-based feature selection

L1 regularization is a technique commonly applied to linear regression and other linear models. It involves adding a regularization term to the loss function to encourage sparsity of model coefficients, thereby enabling feature selection (23).

The goal of feature selection is to choose the most important features from the original feature set to improve model generalization performance and reduce the risk of overfitting. The fundamental principle of L1 regularization is to introduce a penalty term in the loss function, which is the sum of the absolute values of the model coefficients, known as the L1 norm. For linear regression problems, the loss function of L1 regularization can be expressed as:

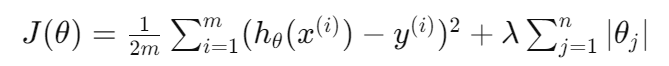

Where:

*J*(*θ*) is the loss function.

*m* is the number of training samples.

*hθ*(*x*(*i*)) is the model's prediction for the *i*-th sample.

*y*(*i*) is the actual output for the *i*-th sample.

*θj* is the *j*-th coefficient of the model.

*N* is the number of features.

*λ* is the regularization parameter controlling the strength of regularization.

The key part of L1 regularization is the term *λ*∑^n^*_j_*_=1_ ∣*θj*∣. This regularization term tends to drive certain feature coefficients to exactly zero, thereby facilitating feature selection. When the coefficient of a feature approaches zero, it indicates that the feature has minimal impact on the model, making it a candidate for exclusion.

By adjusting the regularization parameter *λ*, the sparsity of the coefficients can be controlled to affect the strength of feature selection. Choosing an appropriate *λ* value is a crucial tuning step that typically involves cross-validation on the validation set to find the optimal regularization parameters.

Tree-based feature selection

Tree-based feature selection is a technique that uses decision tree algorithms to assess feature importance and select the most relevant features for a given task (24). Random Forest and Gradient Boosted Trees are commonly used tree-based models for this purpose. Now we will take a deeper look at the principles and methods of tree-based feature selection:

Decision Tree Feature Importance:

Decision trees inherently provide a way to measure the importance of each feature (25). Importance is typically based on how much each feature contributes to reducing impurity (such as, Gini impurity or entropy) across the tree nodes. The more a feature is used to split nodes and reduce impurity, the higher its importance.

Random Forest Feature Importance:

Random Forest builds multiple decision trees and aggregates their predictions(26). Feature importance in a Random Forest is often computed by averaging the feature importance measures across all the trees. The more a feature is used to split nodes and improve the model's performance across various trees, the higher its overall importance.

Gradient Boosted Trees Feature Importance:

Gradient Boosted Trees construct trees sequentially, with the goal of each tree to correct the errors of the previous ones. Feature importance in gradient boosting is calculated by considering how much each feature contributes to the reduction of the loss function (such as mean squared error or cross-entropy) (27). Features that lead to a larger reduction in the loss function have higher importance.

Feature Selection Process:

(1) **Train a Tree-Based Model:** Train a decision tree-based model (Random Forest or Gradient Boosted Trees) on the dataset.

(2) **Calculate** **Feature Importance:** Extract the feature importance scores assigned by the model. This can be done by examining the impurity reduction for each feature in the decision trees or directly using the feature importance measures provided by the model.

(3) **Rank Features:** Rank the features based on their importance scores. The higher the score, the more important the feature is.

(4) **Select Top Features:** Choose the top N features with the highest importance scores. The value of N can be determined based on a predefined threshold or through cross-validation.

Variance threshold feature selection

### The variance of a feature measures the spread or dispersion of its values(28). For a given feature *X*, the variance (*σ*^2^) is calculated as follows:

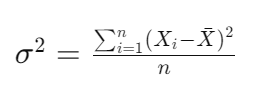

### where:

*n* is the number of samples,

*X_i_* is the value of the feature for the *i*-th sample,

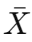
 is the mean of the feature values.

Feature Selection Process:

(1) **Calculate Variances:** Compute the variance of each feature in the dataset.

(2) **Set Threshold:** Define the threshold. Features with variances below this threshold are considered low-variance features.

(3) **Filter Features:** Remove features whose variances are below the threshold. Only retain those features with sufficiently high variance.

**Table S1.** Performance comparison of various feature extraction methods on catBoost

| Feature | Acc | Recall | F1-score | Precision | MCC | AUROC | AUPRC |
| --- | --- | --- | --- | --- | --- | --- | --- |
| All | 0.8769±0.0034 | 0.8896±0.0039 | 0.8902±0.0028 | 0.8907±0.0044 | 0.7677±0.0070 | 0.8640±0.0037 | 0.8671±0.0042 |
| All-AAC | 0.8762±0.0029 | 0.8904±0.0039 | 0.8896±0.0024 | 0.8889±0.0039 | 0.7661±0.0061 | 0.8629±0.0032 | 0.8657±0.0036 |
| All-CKSAAP | 0.8760±0.0025 | 0.8889±0.0041 | 0.8894±0.0021 | 0.8898±0.0040 | 0.7658±0.0052 | 0.8630±0.0028 | 0.8660±0.0033 |
| All-CTDC | 0.8768±0.0030 | 0.8896±0.0045 | 0.9200±0.0026 | 0.9005±0.0034 | 0.7674±0.0063 | 0.8638±0.0031 | 0.8769±0.0033 |
| All-CTDD | 0.8748±0.0020 | 0.8878±0.0039 | 0.8883±0.0015 | 0.8889±0.0049 | 0.7632±0.0043 | 0.8617±0.0027 | 0.8648±0.0036 |
| All-CTDT | 0.8763±0.0029 | 0.8896±0.0053 | 0.8896±0.0026 | 0.8898±0.0038 | 0.7664±0.0060 | 0.8632±0.0030 | 0.8662±0.0033 |
| All-CTriad | 0.8766±0.0021 | 0.8803±0.0039 | 0.8899±0.0018 | 0.8896±0.0038 | 0.7669±0.0044 | 0.8634±0.0024 | 0.8662±0.0030 |
| All-DDE | 0.8772±0.0027 | 0.8900±0.0040 | 0.8804±0.0024 | 0.8908±0.0032 | 0.7683±0.0056 | 0.8642±0.0028 | 0.8673±0.0031 |
| All-DPC | 0.8769±0.0032 | 0.8902±0.0047 | 0.8902±0.0028 | 0.8902±0.0036 | 0.7676±0.0066 | 0.8638±0.0033 | 0.8668±0.0035 |
| All-GAAC | 0.8758±0.0030 | 0.8895±0.0023 | 0.8892±0.0024 | 0.8890±0.0041 | 0.7652±0.0064 | 0.8626±0.0035 | 0.8655±0.0040 |
| All-GDPC | 0.8760±0.0046 | 0.8894±0.0047 | 0.8894±0.0038 | 0.8895±0.0044 | 0.7658±0.0094 | 0.8629±0.0048 | 0.8659±0.0049 |
| All-TPC | 0.8784±0.0024 | 0.8910±0.0045 | 0.8914±0.0019 | 0.8918±0.0050 | 0.7507±0.0050 | 0.8654±0.0030 | 0.8685±0.0038 |
| AAC | 0.8728±0.0033 | 0.8781±0.0032 | 0.8678±0.0027 | 0.8776±0.0041 | 0.7670±0.0069 | 0.8484±0.0037 | 0.8511±0.0039 |
| CKSAAP | 0.8719±0.0026 | 0.8847±0.0041 | 0.8859±0.0022 | 0.8871±0.0048 | 0.7573±0.0056 | 0.8589±0.0032 | 0.8720±0.0039 |
| CTDC | 0.8663±0.0016 | 0.8773±0.0037 | 0.8732±0.0014 | 0.8692±0.0028 | 0.7244±0.0033 | 0.8413±0.0017 | 0.8432±0.0020 |
| CTDD | 0.8028±0.0038 | 0.8381±0.0026 | 0.8251±0.0029 | 0.8124±0.0044 | 0.5892±0.0082 | 0.7722±0.0044 | 0.7787±0.0040 |
| CTDT | 0.8648±0.0040 | 0.8842±0.0010 | 0.8833±0.0031 | 0.8823±0.0058 | 0.7208±0.0087 | 0.8502±0.0049 | 0.8529±0.0053 |
| CTriad | 0.8447±0.0034 | 0.8673±0.0028 | 0.8664±0.0027 | 0.8655±0.0040 | 0.7192±0.0071 | 0.8294±0.0038 | 0.8328±0.0038 |
| DDE | 0.8798±0.0017 | 0.9255±0.0048 | 0.9043±0.0016 | 0.9032±0.0030 | 0.7629±0.0033 | 0.8762±0.0015 | 0.8787±0.0019 |
| DPC | 0.8782±0.0036 | 0.9234±0.0033 | 0.9029±0.0029 | 0.9023±0.0052 | 0.7694±0.0076 | 0.8746±0.0042 | 0.8773±0.0047 |
| GAAC | 0.8136±0.0047 | 0.8384±0.0070 | 0.8327±0.0040 | 0.8271±0.0061 | 0.6329±0.0099 | 0.7854±0.0053 | 0.7914±0.0051 |
| GDPC | 0.8246±0.0064 | 0.8529±0.0047 | 0.8498±0.0052 | 0.8468±0.0067 | 0.6770±0.0136 | 0.8079±0.0070 | 0.8223±0.0068 |
| TPC | 0.8732±0.0022 | 0.8897±0.0047 | 0.8902±0.0020 | 0.8907±0.0035 | 0.7384±0.0045 | 0.8593±0.0023 | 0.8622±0.0026 |
| BEST | 0.8892±0.0046 | 0.9098±0.0045 | 0.9070±0.0033 | 0.9042±0.0093 | 0.7702±0.0102 | 0.8845±0.0062 | 0.8762±0.0074 |

*The bold values indicate the best performance. "BEST" represents a model trained using the remaining descriptors encoding after removing all descriptors with negative impact.

**Table S2.** Performance comparison of various feature extraction methods on XGBoost

| Feature | Acc | Recall | F1-score | Precision | MCC | AUROC | AUPRC |
| --- | --- | --- | --- | --- | --- | --- | --- |
| All | 0.8802±0.0051 | 0.8945±0.0063 | 0.8945±0.0043 | 0.8946±0.0053 | 0.7837±0.0105 | 0.8768±0.0053 | 0.8697±0.0055 |
| All-AAC | 0.8790±0.0023 | 0.8934±0.0014 | 0.8935±0.0018 | 0.8937±0.0037 | 0.7813±0.0049 | 0.8757±0.0028 | 0.8685±0.0033 |
| All-CKSAAP | 0.8816±0.0029 | 0.8946±0.0046 | 0.8956±0.0023 | 0.8966±0.0058 | 0.7866±0.0063 | 0.8785±0.0037 | 0.8716±0.0046 |
| All-CTDC | 0.8785±0.0046 | 0.8914±0.0048 | 0.8930±0.0037 | 0.8946±0.0067 | 0.7803±0.0096 | 0.8754±0.0053 | 0.8687±0.0061 |
| All-CTDD | 0.8790±0.0029 | 0.8930±0.0045 | 0.8935±0.0024 | 0.8939±0.0052 | 0.7812±0.0061 | 0.8757±0.0035 | 0.8686±0.0042 |
| All-CTDT | 0.8806±0.0031 | 0.8941±0.0024 | 0.8948±0.0026 | 0.8956±0.0035 | 0.7847±0.0066 | 0.8775±0.0035 | 0.8705±0.0038 |
| All-CTriad | 0.8804±0.0038 | 0.8947±0.0050 | 0.8947±0.0032 | 0.8947±0.0052 | 0.7840±0.0079 | 0.8770±0.0042 | 0.8698±0.0047 |
| All-DDE | 0.8806±0.0028 | 0.8948±0.0043 | 0.8948±0.0024 | 0.8948±0.0029 | 0.7844±0.0056 | 0.8772±0.0027 | 0.8701±0.0028 |
| All-DPC | 0.8813±0.0042 | 0.8969±0.0052 | 0.8956±0.0035 | 0.8943±0.0049 | 0.7859±0.0086 | 0.8777±0.0044 | 0.8702±0.0048 |
| All-GAAC | 0.8790±0.0015 | 0.8920±0.0047 | 0.8934±0.0011 | 0.8948±0.0053 | 0.7813±0.0034 | 0.8759±0.0025 | 0.8691±0.0036 |
| All-GDPC | 0.8806±0.0026 | 0.8942±0.0067 | 0.8948±0.0022 | 0.8955±0.0057 | 0.7847±0.0053 | 0.8774±0.0030 | 0.8704±0.0039 |
| All-TPC | 0.8798±0.0031 | 0.8948±0.0042 | 0.8943±0.0024 | 0.8938±0.0062 | 0.7829±0.0067 | 0.8763±0.0040 | 0.8690±0.0049 |
| AAC | 0.8709±0.0025 | 0.8868±0.0035 | 0.8867±0.0019 | 0.8867±0.0054 | 0.7944±0.0055 | 0.8672±0.0034 | 0.8899±0.0042 |
| CKSAAP | 0.8656±0.0058 | 0.8857±0.0056 | 0.8826±0.0047 | 0.8795±0.0080 | 0.7831±0.0123 | 0.8609±0.0067 | 0.8830±0.0074 |
| CTDC | 0.8657±0.0042 | 0.8833±0.0044 | 0.8824±0.0036 | 0.8815±0.0035 | 0.7835±0.0086 | 0.8616±0.0042 | 0.8841±0.0042 |
| CTDD | 0.7627±0.0044 | 0.8499±0.0042 | 0.8299±0.0033 | 0.8109±0.0052 | 0.5661±0.0095 | 0.7793±0.0051 | 0.7785±0.0045 |
| CTDT | 0.8543±0.0039 | 0.8741±0.0026 | 0.8729±0.0032 | 0.8717±0.0047 | 0.7597±0.0083 | 0.8796±0.0044 | 0.8723±0.0047 |
| CTriad | 0.8251±0.0044 | 0.8850±0.0048 | 0.8791±0.0035 | 0.8733±0.0055 | 0.6986±0.0093 | 0.8481±0.0050 | 0.8413±0.0050 |
| DDE | 0.8626±0.0041 | 0.8839±0.0046 | 0.8801±0.0033 | 0.8764±0.0052 | 0.7768±0.0085 | 0.8876±0.0045 | 0.8796±0.0048 |
| DPC | 0.8610±0.0031 | 0.8832±0.0030 | 0.8789±0.0022 | 0.8746±0.0064 | 0.7734±0.0069 | 0.8858±0.0043 | 0.8777±0.0052 |
| GAAC | 0.8055±0.0039 | 0.8666±0.0066 | 0.8624±0.0034 | 0.8583±0.0051 | 0.6579±0.0082 | 0.8282±0.0042 | 0.8232±0.0042 |
| GDPC | 0.8163±0.0051 | 0.8728±0.0030 | 0.8711±0.0040 | 0.8695±0.0066 | 0.6809±0.0110 | 0.8401±0.0060 | 0.8346±0.0060 |
| TPC | 0.8255±0.0079 | 0.8529±0.0062 | 0.8791±0.0065 | 0.8753±0.0081 | 0.6996±0.0166 | 0.8491±0.0086 | 0.8425±0.0084 |
| BEST | 0.8854±0.0024 | 0.8701±0.0016 | 0.9005±0.0019 | 0.9009±0.0036 | 0.7837±0.0051 | 0.8819±0.0029 | 0.8748±0.0033 |

### *The bold values indicate the best performance. "BEST" represents a model trained using the remaining descriptors encoding after removing all descriptors with negative impact.

**Table S3.** Performance comparison of various feature extraction methods on RF

| Feature | Acc | Recall | F1-score | Precision | MCC | AUROC | AUPRC |
| --- | --- | --- | --- | --- | --- | --- | --- |
| All | 0.6112±0.0114 | 0.9962±0.0025 | 0.7531±0.0051 | 0.6054±0.0073 | 0.1342±0.0552 | 0.5209±0.0147 | 0.6053±0.0073 |
| All-AAC | 0.6055±0.0025 | 0.9975±0.0013 | 0.7506±0.0010 | 0.6016±0.0016 | 0.1153±0.0132 | 0.5136±0.0033 | 0.6016±0.0016 |
| All-CKSAAP | 0.6022±0.0027 | 0.9986±0.0003 | 0.7492±0.0012 | 0.5995±0.0016 | 0.0939±0.0223 | 0.5092±0.0034 | 0.5995±0.0016 |
| All-CTDC | 0.6174±0.0100 | 0.9956±0.0029 | 0.7559±0.0044 | 0.6093±0.0067 | 0.1684±0.0360 | 0.5286±0.0131 | 0.6092±0.0066 |
| All-CTDD | 0.6082±0.0083 | 0.9966±0.0022 | 0.7517±0.0036 | 0.6034±0.0053 | 0.1221±0.0434 | 0.5171±0.0107 | 0.6034±0.0053 |
| All-CTDT | 0.6117±0.0087 | 0.9963±0.0015 | 0.7533±0.0039 | 0.6056±0.0055 | 0.1408±0.0429 | 0.5214±0.0110 | 0.6056±0.0055 |
| All-CTriad | 0.6146±0.0103 | 0.9936±0.0040 | 0.7542±0.0043 | 0.6078±0.0070 | 0.1501±0.0419 | 0.5257±0.0136 | 0.6077±0.0069 |
| All-DDE | 0.6075±0.0102 | 0.9977±0.0026 | 0.7516±0.0045 | 0.6029±0.0065 | 0.1147±0.0587 | 0.5159±0.0131 | 0.6028±0.0064 |
| All-DPC | 0.6055±0.0032 | 0.9972±0.0018 | 0.7505±0.0012 | 0.6017±0.0022 | 0.1139±0.0145 | 0.5136±0.0043 | 0.6016±0.0022 |
| All-GAAC | 0.6076±0.0029 | 0.9987±0.0008 | 0.7518±0.0014 | 0.6027±0.0018 | 0.1306±0.0185 | 0.5158±0.0036 | 0.6027±0.0018 |
| All-GDPC | 0.6097±0.0061 | 0.9965±0.0016 | 0.7524±0.0027 | 0.6044±0.0038 | 0.1333±0.0349 | 0.5190±0.0077 | 0.6043±0.0038 |
| All-TPC | 0.6823±0.0091 | 0.9764±0.0043 | 0.7853±0.0046 | 0.6568±0.0074 | 0.3481±0.0204 | 0.6133±0.0118 | 0.6554±0.0071 |
| AAC | 0.8153±0.0091 | 0.9215±0.0081 | 0.8559±0.0056 | 0.7993±0.0130 | 0.6148±0.0184 | 0.7905±0.0123 | 0.7832±0.0109 |
| CKSAAP | 0.6769±0.0175 | 0.9698±0.0094 | 0.7814±0.0079 | 0.6545±0.0147 | 0.3284±0.0401 | 0.6081±0.0235 | 0.6526±0.0138 |
| CTDC | 0.8036±0.0125 | 0.8910±0.0089 | 0.8437±0.0095 | 0.8013±0.0115 | 0.5875±0.0270 | 0.7831±0.0138 | 0.7789±0.0115 |
| CTDD | 0.6392±0.0126 | 0.9597±0.0168 | 0.7600±0.0038 | 0.6293±0.0111 | 0.2161±0.0342 | 0.5640±0.0190 | 0.6278±0.0102 |
| CTDT | 0.6052±0.0024 | 0.9976±0.0013 | 0.7504±0.0009 | 0.6014±0.0016 | 0.1133±0.0123 | 0.5131±0.0033 | 0.6014±0.0016 |
| CTriad | 0.6558±0.0105 | 0.9819±0.0017 | 0.7725±0.0052 | 0.6368±0.0073 | 0.2822±0.0304 | 0.5793±0.0131 | 0.6360±0.0071 |
| DDE | 0.6677±0.0086 | 0.9707±0.0046 | 0.7766±0.0045 | 0.6472±0.0064 | 0.3059±0.0224 | 0.5966±0.0107 | 0.6457±0.0061 |
| DPC | 0.6799±0.0127 | 0.9712±0.0070 | 0.7831±0.0061 | 0.6562±0.0104 | 0.3376±0.0292 | 0.6115±0.0166 | 0.6544±0.0098 |
| GAAC | 0.8305±0.0053 | 0.8475±0.0067 | 0.8561±0.0039 | 0.8651±0.0096 | 0.6504±0.0119 | 0.8266±0.0068 | 0.8239±0.0073 |
| GDPC | 0.8064±0.0022 | 0.8867±0.0078 | 0.8449±0.0024 | 0.8070±0.0030 | 0.5935±0.0048 | 0.7875±0.0019 | 0.7829±0.0017 |
| TPC | 0.5983±0.0027 | 0.9993±0.0007 | 0.7475±0.0012 | 0.5970±0.0017 | 0.0616±0.0223 | 0.5042±0.0034 | 0.5970±0.0017 |
| BEST | 0.8416±0.0034 | 0.9073±0.0032 | 0.8720±0.0021 | 0.8394±0.0058 | 0.6684±0.0071 | 0.8261±0.0047 | 0.8168±0.0047 |

### *The bold values indicate the best performance. "BEST" represents a model trained using the remaining descriptors encoding after removing all descriptors with negative impact.

### Table S4. Performance evaluation of different feature selection methods on 5-fold cross-validation.

| Method | Acc | Recall | F1-score | Precision | MCC | AUROC | AUPRC |
| --- | --- | --- | --- | --- | --- | --- | --- |
| chi2 | 0.8814± 0.0030 | 0.9015±0.0074 | 0.9002±0.0024 | 0.8991±0.0071 | 0.7542±0.0065 | 0.8768±0.0039 | 0.8689±0.0049 |
| Basd-L1 | 0.6669±0.0184 | 0.8077±0.0156 | 0.7422±0.0134 | 0.6868±0.0163 | 0.2878±0.0425 | 0.6345±0.0211 | 0.6689±0.0134 |
| SelectFromRF | 0.8858±0.0036 | 0.9082±0.0043 | 0.9042±0.0027 | 0.9003±0.0067 | 0.7629±0.0078 | 0.8806±0.0046 | 0.8721±0.0054 |
| VarianceThreshold | 0.8867±0.0035 | 0.9054±0.0046 | 0.9046±0.0027 | 0.9039±0.0071 | 0.7651±0.0078 | 0.8824±0.0047 | 0.8745±0.0056 |

*The bold values indicate the best performance.

### Table S5. Performance evaluation of different feature selection methods on the independent test.

| Method | Acc | Recall | F1-score | Precision | MCC | AUROC | AUPRC |
| --- | --- | --- | --- | --- | --- | --- | --- |
| chi2 | 0.8442 | 0.88 | 0.8502 | 0.8224 | 0.69 | 0.844 | 0.784 |
| Basd-L1 | 0.6331 | 0.8 | 0.6867 | 0.6015 | 0.281 | 0.6323 | 0.5817 |
| SelectFromRF | 0.8392 | 0.84 | 0.8285 | 0.8571 | 0.6787 | 0.8392 | 0.8 |
| VarianceThreshold | 0.799 | 0.85 | 0.8095 | 0.7727 | 0.6008 | 0.7987 | 0.7322 |

*The bold values indicate the best performance.

### Table S6. Performance comparison of the whole sequence models without FS, with FS, and with FS + SMOTE on 5-fold cross-validation.

| Method | Acc | Recall | F1-score | Precision | MCC | AUROC | AUPRC |
| --- | --- | --- | --- | --- | --- | --- | --- |
| WSM (w/o FS) | 0.8892±0.0046 | 0.9098±0.0045 | 0.9070±0.0033 | 0.9042±0.0093 | 0.7702±0.0102 | 0.8845±0.0062 | 0.8762±0.0074 |
| WSM (With FS) | 0.8814± 0.0030 | 0.9015±0.0074 | 0.9002±0.0024 | 0.8991±0.0071 | 0.7542±0.0065 | 0.8768±0.0039 | 0.8689±0.0049 |
| WSM (FS+SMOTE) | 0.8817±0.0037 | 0.8999±0.0054 | 0.9003±0.0025 | 0.9008±0.0091 | 0.7549±0.0085 | 0.8775±0.0055 | 0.8700±0.0068 |

*The bold values indicate the best performance. “WSM” the means whole sequence model “FS” means feature selection. The performance was evaluated based on the catBoost model with the screened features.

### Table S7. Performance comparison of whole sequence-based models with different processing methods on the independent test.

| Method | Acc | Recall | F1-score | Precision | MCC | AUROC | AUPRC |
| --- | --- | --- | --- | --- | --- | --- | --- |
| WSM (w/o FS) | 0.82 | 0.8 | 0.8163 | 0.8333 | 0.6405 | 0.82 | 0.7667 |
| WSM (With FS) | 0.845 | 0.88 | 0.8502 | 0.8224 | 0.6917 | 0.845 | 0.7837 |
| WSM (FS+SMOTE) | 0.82 | 0.83 | 0.8218 | 0.8137 | 0.6401 | 0.82 | 0.7604 |

*The bold values indicate the best performance. “WSM” means whole sequence model. “FS” means feature selection. The performance was evaluated based on the catBoost model with screened features.

### References

1.	Chen Z, Zhao P, Li F, Leier A, Marquez-Lago TT, Wang Y, et al. iFeature: a python package and web server for features extraction and selection from protein and peptide sequences. Bioinformatics. 2018;34(14):2499-502.

2.	Li F, Leier A, Liu Q, Wang Y, Xiang D, Akutsu T, et al. Procleave: Predicting Protease-specific Substrate Cleavage Sites by Combining Sequence and Structural Information. Genomics Proteomics Bioinformatics. 2020;18(1):52-64.

3.	Song J, Wang Y, Li F, Akutsu T, Rawlings ND, Webb GI, et al. iProt-Sub: a comprehensive package for accurately mapping and predicting protease-specific substrates and cleavage sites. Brief Bioinform. 2019;20(2):638-58.

4.	Li F, Zhang Y, Purcell AW, Webb GI, Chou KC, Lithgow T, et al. Positive-unlabelled learning of glycosylation sites in the human proteome. BMC Bioinf. 2019;20(1):112.

5.	Chen K, Kurgan LA, Ruan J. Prediction of flexible/rigid regions from protein sequences using k-spaced amino acid pairs. BMC Struct. Biol. 2007;7(1):1-13.

6.	Usman M, Lee JA, editors. Afp-cksaap: Prediction of antifreeze proteins using composition of k-spaced amino acid pairs with deep neural network. 2019 IEEE 19th International Conference on Bioinformatics and Bioengineering (BIBE); 2019: IEEE.

7.	Wang Y, Li F, Bharathwaj M, Rosas NC, Leier A, Akutsu T, et al. DeepBL: a deep learning-based approach for in silico discovery of beta-lactamases. Brief Bioinform. 2020.

8.	Li F, Li C, Wang M, Webb GI, Zhang Y, Whisstock JC, et al. GlycoMine: a machine learning-based approach for predicting N-, C- and O-linked glycosylation in the human proteome. Bioinformatics. 2015;31(9):1411-9.

9.	Li F, Li C, Revote J, Zhang Y, Webb GI, Li J, et al. GlycoMine(struct): a new bioinformatics tool for highly accurate mapping of the human N-linked and O-linked glycoproteomes by incorporating structural features. Sci Rep. 2016;6:34595.

10.	Chen Z, Zhao P, Li C, Li F, Xiang D, Chen Y-Z, et al. iLearnPlus: a comprehensive and automated machine-learning platform for nucleic acid and protein sequence analysis, prediction and visualization. Nucleic Acids Res. 2021;49(10):e60-e.

11.	Saravanan V, Gautham N. Harnessing computational biology for exact linear B-cell epitope prediction: a novel amino acid composition-based feature descriptor. OMICS. 2015;19(10):648-58.

12.	Bhasin M, Raghava G. ESLpred: SVM-based method for subcellular localization of eukaryotic proteins using dipeptide composition and PSI-BLAST. Nucleic Acids Res. 2004;32(suppl_2):W414-W9.

13.	Charoenkwan P, Nantasenamat C, Hasan MM, Moni MA, Manavalan B, Shoombuatong W. UMPred-FRL: A new approach for accurate prediction of umami peptides using feature representation learning. Int. J. Mol. Sci. 2021;22(23):13124.

14.	Dholaniya PS, Rizvi S. Effect of various sequence descriptors in predicting human proteinprotein interactions using ANN-based prediction models. Curr Bioinform. 2021;16(8):1024-33.

15.	Elumalai E, Muthuvel SK. Characterization and Prediction of Dengue Virus Targeting Peptides Based on Combined Amino Acid Composition Descriptors Using Random Forest Algorithm. 2021.

16.	Basith S, Lee G, Manavalan B. STALLION: a stacking-based ensemble learning framework for prokaryotic lysine acetylation site prediction. Brief Bioinform. 2022;23(1):bbab376.

17.	Prokhorenkova L, Gusev G, Vorobev A, Dorogush AV, Gulin A. CatBoost: unbiased boosting with categorical features. NeurIPS. 2018;31.

18.	Hancock JT, Khoshgoftaar TM. CatBoost for big data: an interdisciplinary review. Journal of big data. 2020;7(1):1-45.

19.	Chen T, Guestrin C. XGBoost: A Scalable Tree Boosting System. Proceedings of the 22nd ACM SIGKDD International Conference on Knowledge Discovery and Data Mining; San Francisco, California, USA: Association for Computing Machinery; 2016. p. 785–94.

20.	Breiman L. Random forests. Mach Learn. 2001;45:5-32.

21.	I KABACOFF R. Data analysis and graphics with R. 2011.

22.	Peker N, Kubat C. Application of Chi-square discretization algorithms to ensemble classification methods. Expert Syst. Appl. 2021;185:115540.

23.	Schmidt M, Fung G, Rosales R. Optimization methods for l1-regularization. University of British Columbia, Technical Report TR-2009-19. 2009.

24.	Zhou H, Zhang J, Zhou Y, Guo X, Ma Y. A feature selection algorithm of decision tree based on feature weight. Expert Syst. Appl. 2021;164:113842.

25.	Al Iqbal MR, Rahman S, Nabil SI, Chowdhury IUA, editors. Knowledge based decision tree construction with feature importance domain knowledge. 2012 7th international conference on electrical and computer engineering; 2012: IEEE.

26.	Menze BH, Kelm BM, Masuch R, Himmelreich U, Bachert P, Petrich W, et al. A comparison of random forest and its Gini importance with standard chemometric methods for the feature selection and classification of spectral data. BMC Bioinf. 2009;10:1-16.

27.	Adler AI, Painsky A. Feature importance in gradient boosting trees with cross-validation feature selection. Entropy. 2022;24(5):687.

28.	Fida MAFA, Ahmad T, Ntahobari M, editors. Variance threshold as early screening to Boruta feature selection for intrusion detection system. 2021 13th International Conference on Information & Communication Technology and System (ICTS); 2021: IEEE.
